# Multistate Ornstein-Uhlenbeck approach for practical estimation of movement and resource selection around central places

**DOI:** 10.1101/2020.10.28.359430

**Authors:** Joseph M. Eisaguirre, Travis L. Booms, Christopher P. Barger, Scott D. Goddard, Greg A. Breed

## Abstract

1. Home range dynamics and movement are central to a species’ ecology and strongly mediate both intra- and interspecific interactions. Numerous methods have been introduced to describe animal home ranges, but most lack predictive ability and cannot capture effects of dynamic environmental patterns, such as the impacts of air and water flow on movement.
2. Here, we develop a practical, multi-stage approach for statistical inference into the behavioral mechanisms underlying how habitat and dynamic energy landscapes—in this case how airflow increases or decreases the energetic efficiency of flight—shape animal home ranges based around central places. We validated the new approach using simulations, then applied it to a sample of 12 adult golden eagles *Aquila chrysaetos* tracked with satellite telemetry.
3. The application to golden eagles revealed effects of habitat variables that align with predicted behavioral ecology. Further, we found that males and females partition their home ranges dynamically based on uplift. Specifically, changes in wind and sun angle drove differential space use between sexes, especially later in the breeding season when energetic demands of growing nestlings require both parents to forage more widely.
4. This method is easily implemented using widely available programming languages and is based on a hierarchical multistate Ornstein-Uhlenbeck space use process that incorporates habitat and energy landscapes. The underlying mathematical properties of the model allow straightforward computation of predicted utilization distributions, permitting estimation of home range size and visualization of space use patterns under varying conditions.

## Introduction

The “home range” has been a central concept in animal behavior for some time (Burt, 1943; Dunn and Gipson, 1977). To measure and understand an animal’s home range—the area in which an animal carries out its regular foraging and reproductive activities (Burt, 1943)—researchers have applied techniques ranging from simple and purely descriptive, such as methods like minimum convex polygons and kernel density estimators, to complex mechanistic models, such as advection-diffusion equations (Moorcroft and Lewis, 2006; Hooten et al., 2017). Along this spectrum of complexity are a set of analyses of intermediate complexity known as resource selection functions (RSFs; Manly et al., 2002) and related step selection functions (SSFs; Fortin et al., 2005). The RSF and SSF frameworks separate the probability of an animal occurring at a location on the landscape into two parts: availability (or movement) and resource selection (Moorcroft and Barnett, 2008). Together, movement and resource weighting functions can describe an array of animal space use patterns (Potts et al., 2014b).

One early conceptual model of animal space use dynamics was the “elastic disc hypothesis,” which describes animal space use as the degree to which boundaries of territories are compressible, shaped by the territorial aggression of neighboring conspecifics (Huxley, 1934). This process is analogous to the way an elastic disc can be molded by extrinsic forces, and the analogy forms a general conceptual foundation describing the formation and dynamics of animal home ranges (Getty, 1981). For example, consider an animal that requires a certain amount of suitable habitat. Given no extrinsic forces, that animal might spend much of its time within a smaller core area, venturing out equally in all directions to acquire resources. This would give rise to a circular or disc-shaped home range, and would be especially true for an animal that has a “central place” such as a nest or den that requires tending. In contrast, where an animal resides near the boundary of suitable habitat, its home range must stretch along that boundary, as the amount of suitable habitat required remains constant, and the shape of the home range will consequently conform to habitat constraints.

In reality, habitat constraints can change through time. However, many of the more common approaches to quantifying animal home ranges describe animal space use as static in time, either because the descriptive method cannot accommodate time or home ranges are actually assumed to be static. Animal movement, however, is usually much more fluid, driven by suites of intrinsic and extrinsic forces (Nathan et al., 2008). Consequently, home ranges are fundamentally dynamic.

Forces that drive these dynamics include the energy landscape, a conceptual framework that incorporates how an animal’s movement can be shaped by its energetic demands interacting with dynamic landscape features, especially moving fluids such as air or water (Shepard et al., 2013). These dynamics alter a landscape’s suitability and shape space use patterns in a number of ways (Morales and Ellner, 2002; Schooley and Wiens, 2004; Prokopenko et al., 2016). For animals that can take advantage of variable energy subsidies available from moving fluids, including soaring birds that use uplift and aquatic animals that ride water currents, dynamic space use patterns and emergent home range properties will be shaped by these features (Shepard et al., 2013). In such situations, the elastic disc will constantly vary, changing shape as the weather changes.

The RSFs and SSFs noted earlier are widely used and generally robust quantitative assessments of animal space use and home range dynamics, and they have been continuously refined and improved since their respective introductions. Getty (1981) presented an early RSF adaptation inspired by the elastic disc hypothesis. Another early model has also been considered in understanding animal home ranges—the Ornstein-Uhlenbeck (OU) proccess (Dunn and Gipson, 1977)—and can relate to the elastic disc hypothesis.

Here, we develop a practical hierarchical modelling approach for inferring the mechanisms of home range dynamics and how habitat and the energy landscape interact with behavior to shape animal home ranges. This method combines the OU and SSF modelling frameworks and operates on landscapes with dynamic energy subsidies driven by atmospheric forcing. The approach is similar to but extends some previously introduced methods (i.e. Johnson et al., 2008; Christ et al., 2008), which have not seemed to gain traction among practitioners, likely due to computational limitations and difficulty in application. Our approach overcomes computational issues and eases application without sacrificing inference, which we validated with a simulation study. Finally, we applied it to analyze the home range behavior and space use of territorial golden eagles *Aquila chrysaetos*. Specifically, we fit models to estimate how male and female territorial eagles partitioned space during the breeding season based on different habitats or dynamic features of the landscape (i.e. thermal and orographic uplift).

## Methods

### Ornstein-Uhlenbeck home range model

An OU process over two-dimensional space is continuous-time, mean-reverting, and can help researchers study home range behavior of animals that tend a central place (e.g., a nest; Dunn and Gipson, 1977; Blackwell, 1997; Breed et al., 2017). Assuming independence in the two spatial dimensions simplifies the model and aligns better with central place behavior, as movement is equally likely in all directions around the central point. Such an OU process can be presented as the following stochastic differential equation (SDE):

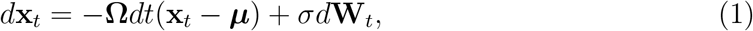

where **x**_*t*_ is a coordinate vector of the location of the animal at time *t*, **Ω**= *ω***I**_2_ with *ω* describing the strength of the animal’s tendency to move toward the central point ***μ***, *σ >* 0, and **W**_*t*_ is Brownian motion. The solution of this SDE takes the form:

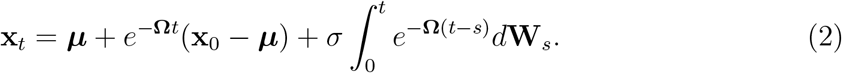

While this solution conveniently gives the position of the animal at any time *t*, we typically observe animal movement by recording series of discrete locations by, for example, using radio or GPS telemetry. This invokes the position likelihood of the OU process:

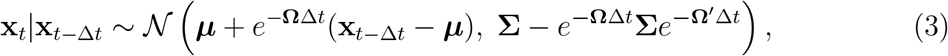

where **Σ**= *σ*^2^**I**_2_. This discretized formulation can be described as a biased random walk (BRW) with a bias toward ***μ***. Notably, it reaches a long term steady state 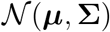 due to the rapidly decaying effect of conditioning on **x**_*t*_ as Δ*t* increases (Blackwell, 1997).

Assuming independence in the two spatial dimensions helps wed the OU process to the elastic disc hypothesis (Huxley, 1934; Getty, 1981), similar to the circular normal distribution used by Getty (1981). A chosen contour of 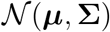 can be a circular approximation of an animal’s home range. Further, the highest probability density value of 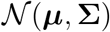 is centered on ***μ***, consistent with central place behavior. Note that using equation (8) takes into account serial correlation, which is inherent to an animal’s movement, ensuring an unbiased estimate of **Σ**. Additionally, the continuous-time nature of the process makes it applicable under any temporal resolution of data and any irregularities in that data.

The shape of the home range may be modified by various extrinsic factors (Getty, 1981), which can be built into the OU process with an RSF in the weighted distribution framework (Johnson et al., 2008). The general form of this framework describes the probability density *f_u_* of an animal’s location over some landscape **z** containing a suite of habitat types and resources as the product of a density explaining what is available to the animal *f_a_* and a weighting function *ψ*:

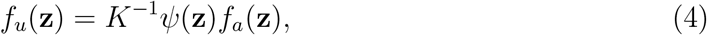

where *K* is a normalizing constant. When *f_a_* takes the form of an OU process (equation 8) and *ψ*(**z**(**x**_*t*_)) = exp[**z**(**x**_*t*_)′**β**], where the function **z**(**x**_*t*_) returns a vector of habitat values and/or resources associated with a location **x**_*t*_ that lies in **z** and **β** weights those resources based on the animal’s preferences, the conditional probability density of the location of the animal can be written as

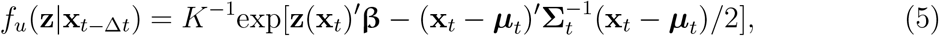

where ***μ****_t_* = ***μ*** + *e^**−**^*^**Ω**Δ*t*^(**x**_*t*−Δ*t*_ − ***μ***) and **Σ**_*t*_ = **Σ**− *e^**−**^*^**Ω**Δ*t*^**Σ***e*^−**Ω**^1^Δ*t*^ (Johnson et al., 2008). Note that the habitat covariates (**z**(**x**_*t*_)) are spatiotemporally explicit so that the effects of dynamic habitat and landscape variables may be accounted for in estimation of parameters and predicting utilization distributions.

#### Multi-stage estimation

Evaluating *K* is usually problematic but often avoided in estimating **β**, as with more conventional RSF models, by implementing an use-availability design that compares resources at ‘available’ locations to ‘used’ locations with logistic regression (Lele and Keim, 2006; Hooten et al., 2017). We note that equation (5) resembles a more conventional RSF model with an offset term—the anisotropic distance between **x**_*t*_ and **x**_*t*−Δ*t*_ (Johnson et al., 2008). We consequently posited that if the OU process parameters were estimated first, then were used to construct the necessary covariate (i.e. 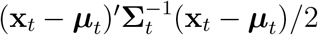, **β** could then be estimated in a second step with regression, which is similar to constructing covariates for estimating **β** with Poisson regression (Johnson et al., 2013) and conditional logistic regression (Forester et al., 2009). Although a sacrifice in statistical elegance, this saves considerable model complexity and estimation challenge, especially when hierarchical inference of **β** across several individuals is a primary goal. As we show, the inference achieved with this procedure does not meaningfully differ from the more elegant, but far more difficult approach, described by Johnson et al. (2008), and makes available hierarchical estimation that is not possible with their method.

Our proposed estimation procedure is as follows. First, estimate the movement parameters in equation 8. Second, use those fitted parameters to make predictions about each **x**_*t*_, effectively generating so-called available locations. Assuming estimation of equation 8 is done in a Bayesian framework, this second step involves sampling from the marginal posterior predictive distributions (Hooten et al., 2014, 2017; Eisaguirre et al., 2020). Next, the quantities 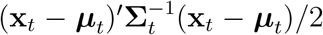 are computed using point estimates of parameters. Finally, the selection coefficients **β** are estimated using logistic regression, as in conventional use-availability resource selection analysis designs, which, in the Bayesian framework, is an empirical Bayes procedure.

If multilevel inference across several individual animals is desired, it is typically straightforward to incorporate such complexity in the regression model for estimating **β**. Higher level inference of the movement parameters may pose a challenge, however, in which case recursive Bayesian inference could be used (Lunn et al., 2013; Hooten et al., 2016; Hooten and Hefley, 2019); we detail this in the Model Extensions section below. Of course, such could be used for estimating **β** as well, if the regression model structure poses estimation challenges.

#### Dynamic utilization distributions

An advantage of the OU model within this framework is that it explicitly weights locations closer to the central point ***μ*** more heavily. If it did not, space use in that area would be attributed solely to habitat or resources there, as opposed to availability, which could bias 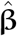. Another advantage of this OU model is that it can be used to compute home range estimates from a set of hypothesized mechanisms, such as different, possibly interacting, and/or dynamic habitat variables. Given that *ψ* is assumed stationary and as Δ*t* gets large *f_a_* approaches 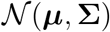,

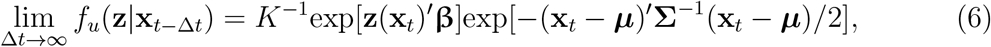

which is simply the normalized product of a multivariate normal kernel and the habitat weighting function. We are thus left with habitat-independent central place (circular) home range estimator 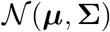 and a weighting function *ψ*(**z**) = exp[**z**(**x**_*t*_)′**β**] that shapes the home range (equation 6). The product of these provides the stationary estimate of *f_u_*, a contour of which is the mathematical description of the conceptual elastic disc (Huxley, 1934) molded by the habitat (Fig. 1). Further, when the resources over the landscape **z** vary through time and are dynamic, evaluating the steady state of *f_u_* must be done with resource values **z**(**x**_*t*_*) fixed at some hypothetical or characteristic time *t* = *t*^*^. We can thus choose *t*^*^ to make predictions about how space use changes based on dynamic resources. This is in contrast to many RSF and SSF studies in the literature, which are typically restricted to evaluating *ψ*(**z**(**x**_*t*_*)), rather than the utilization distribution *f_u_*(**z**(**x**_*t*_*)).

**Figure 1:**
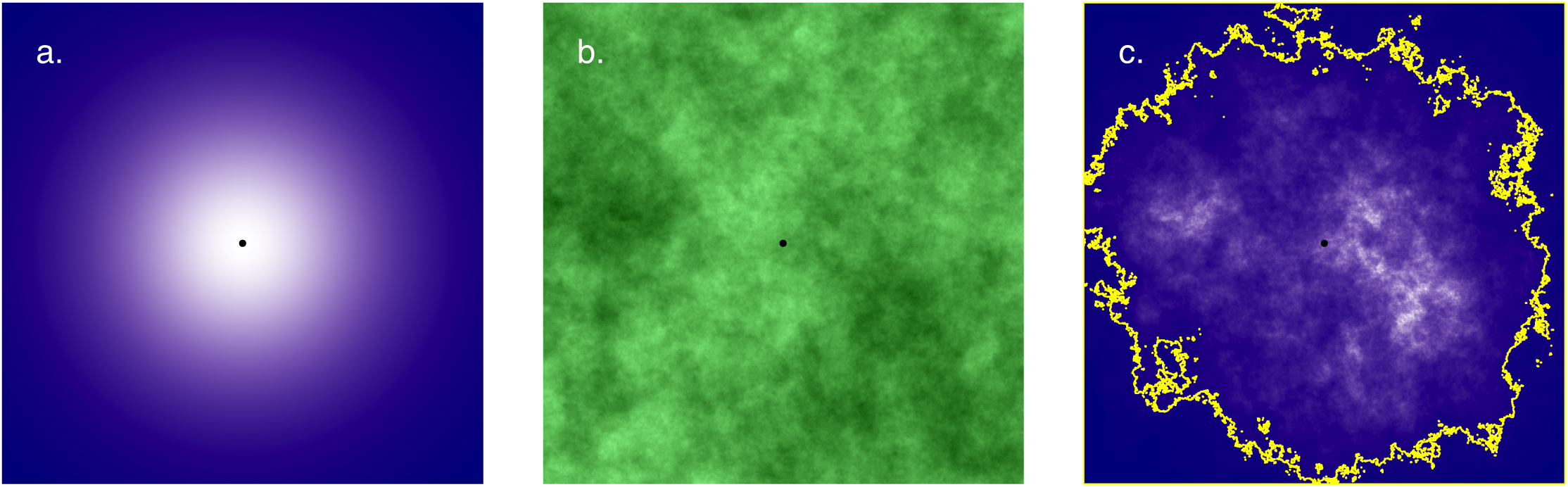
Example of computing the steady-state, analytical home range and space use distribution from an Ornstein-Uhlenbeck space use model. The movement-only, habitat-independent space use distribution (a) is modified by the habitat (b) and the animal’s preferences for that habitat (i.e. a habitat weighting function), giving rise to a predicted space use distribution (c). Point is animal’s center of attraction, and the polygon in c is the 95% volume contour of the space use distribution, representing an estimated home range boundary.

### Simulation study

#### Methods

To ensure that estimation of the OU process and resource selection parameter estimates were unbiased and informative when estimated with the multi-stage procedure, we conducted a simulation study generally following the approaches of Forester et al. (2009) and Johnson et al. (2008). The simulation began with the creation of three artificial landscapes containing a continuous resource variable. Using R and the package `RandomFields` (R Core Team, 2018; Schlather et al., 2019), landscapes were generated on a 2000 × 2000 grid using a Gaussian random field (GRF) with an exponential covariance function. The scale parameter was set at 10, 50, or 100, prescribing each landscape a different level of spatial autocorrelation. We simulated 100 tracks, each 100 movements in length, for each landscape and each of six parameter combinations (*β* = 0, 1, or 2 and *ω* = 1 or 2) for a total of 18 landscape/parameter scenarios. *σ*^2^ was fixed at 100^2^ and ***μ*** at (1000, 1000). Additionally, to ensure identifibility of **β** in the case of multiple covariates, we did one simulation with the scale parameter set to 100, *ω* = 2, *β*_1_ = 1, and *β*_2_ = 1, where *β*_2_ is the coefficient for a binary covariate covering half of the spatial domain. For each simulated track, we fit the OU model, assuming the central point ***μ*** known, generated available points, computed the necessary covariate from the estimated OU parameters, and then attempted to estimate *β* with an use-availability design using logistic regression.

Estimation was performed in a Bayesian framework using Stan and R (Stan Development Team, 2016, 2018; R Core Team, 2018), sampling five available points for each used point from the marginal posterior predictive distributions of each **x**_*t*_ (Hooten et al., 2014, 2017; Eisaguirre et al., 2020). We used three chains of 15,000 Hamiltonian Monte Carlo (HMC) iterations, including 5,000 for warm-up, and retained 1,000 samples for inference in fitting the OU movement model. We used four chains of 5,000 iterations, including 3,000 for warmup, and retained 2,000 samples for inference in estimating the selection parameter *β*. Weakly informative (truncated) normal priors were placed on the OU parameters, centered away from the true values, and a weakly informative normal prior on *β*, centered on zero. See Appendix 2 for code containing details about the priors. The covariate that accounts for the OU movement process in estimating *β* (i.e. 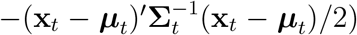 was computed for each used and available point using the posterior means of *ω* and *σ*^2^. *β* was then estimated with an use-availability design and Bayesian logistic regression. For each parameter combination, we summarized the relative biases of the posterior means and the proportion of tracks for which the 95% credible interval overlapped the true value for *β*, *ω*, and *σ*^2^.

#### Simulation Results

The proportions of 95% credible interval coverage were *>* 0.80 for nearly all cases in estimates of *β* (three were *>* 0.70) and generally high for *σ*^2^ and *ω* as well (Figs. S1 & S2). The simulation to assure identifiability found high credible interval coverage (*>* 0.80) as well. Thus, simulations generally found the two-step approach provided estimates of resource selection parameters *β* with no or minimal bias (Fig. 2). Other use-availability designs have also been found to yield unbiased estimates of resource selection parameters (Lele and Keim, 2006; Forester et al., 2009; Avgar et al., 2016). Estimating the movement parameters *ω* and *σ*^2^ yielded slightly more bias, but Johnson et al. (2008) had similar levels of bias when maximizing the joint likelihood for equation (5) rather than the simpler two-step procedure we describe.

**Figure 2:**
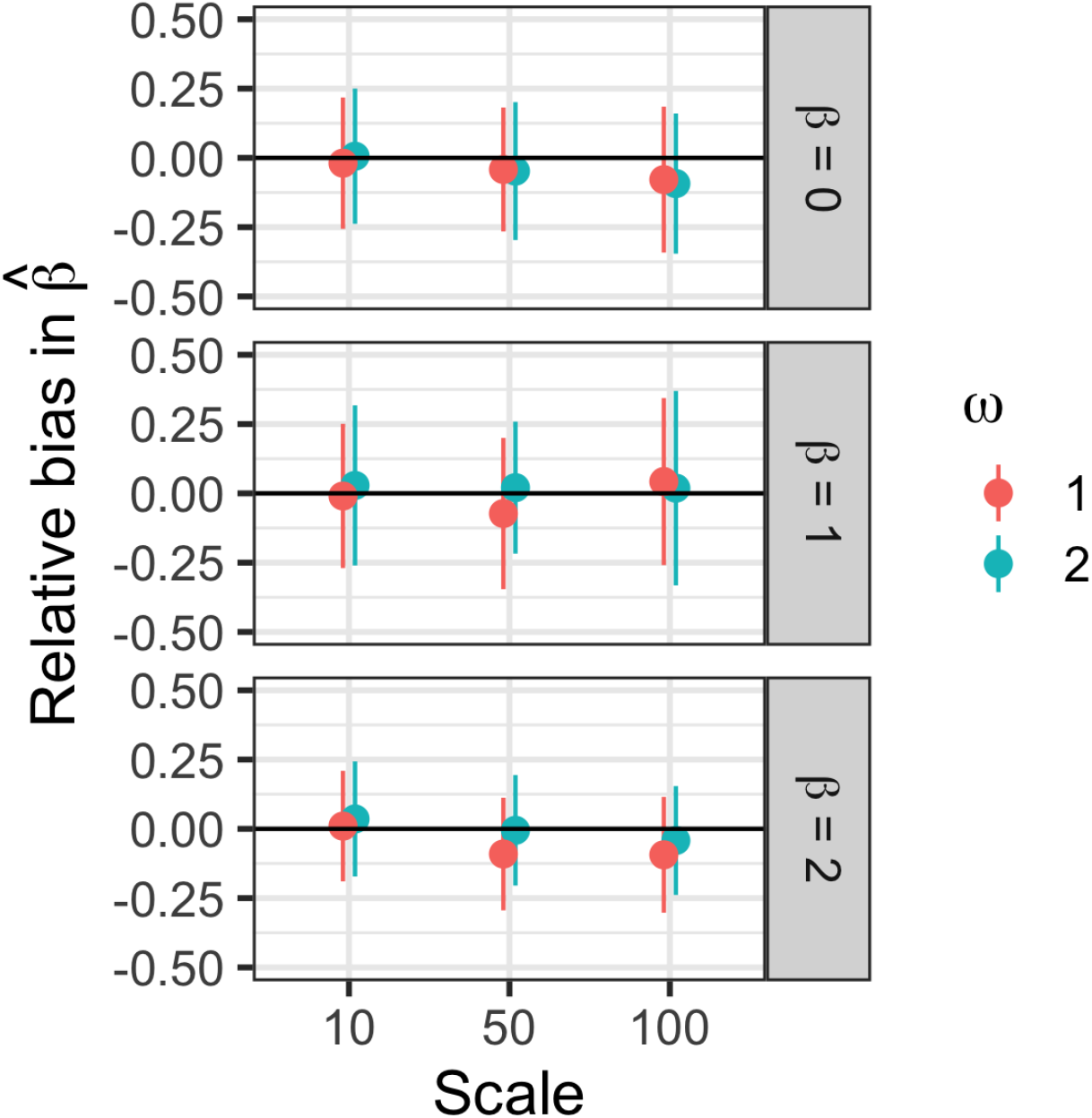
Summary of the relative bias in the selection coefficient *β* when estimated with an Ornstein-Uhlenbeck home range model with movement parameters estimated offline. The ‘scale’ parameter adjusts the level of spatial autocorrelation over the artificial landscape movements were simulated on, and *ω* is a movement parameter. Points are the average of posterior means computed across 100 simulations ± two standard deviations.

### Model extensions

#### Multiple home range cores

An OU home range model can be extended to allow for multiple core areas, and each core can be allowed to have a unique set of movement patterns within an animal’s broader home range (Johnson et al., 2008; Breed et al., 2017). One way to accomplish this is estimating transitions among *K* cores as a Markov process, with a *K* × *K* transition matrix **Γ** describing the probability of the animal moving from one core to another (or remaining in the currently occupied core) during the time interval *t* to *t* + 1 (Breed et al., 2017). Note that to ensure the Markov assumptions hold, fixed and regular time intervals are required, which is common in most (but not all) types of telemetry data. If data are not regular, one can simply use an indexing approach to still incorporate multiple cores (*sensu* Johnson et al., 2008). We can also estimate the relationships between transition probabilities and habitat conditions or other covariates in a manner similar to multinomial logistic regression. Breed et al. (2017) estimated parameters associated with staying in a core area (i.e. the elements along the diagonal of **Γ**); however, here we extend that to transitions among all cores. As these covariates can be temporally dynamic, we may denote our transition matrix as **Γ**_*t*_ = (*γ_ij,t_*). Employing the multinomial logit link, we can write the conditional probability that the animal is in the *j*th core at time *t* + 1 given that it came from the *i*th core:

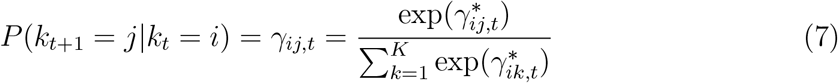

where 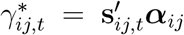 is the vector of covariates associated with the core *k_t_* = *i* at time *t*, and the vector ***α***_*ij*_ weights those covariates by their effect on *γ_ij,t_*. We could thus calculate **Γ**_*t*_ for a set of core- and time-specific covariates. This is similar to modeling behavioral state transitions with a conventional hidden Markov Model for animal movement data (*sensu* Michelot et al., 2016), but the ‘states’ here are home range cores, each having a respective set of movement parameters (Breed et al., 2017).

Unsupervised estimation of the state transitions, which in Stan required marginalizing the latent discrete process, proved computationally impractical. We thus followed Breed et al. (2017) and implemented a *k*-means clustering algorithm to identify the number of home range core areas, the location of each core center ***μ_k_***, and the core transitions a priori (Hartigan and Wong, 1979). We then proceeded with supervised estimation of ***α*** and assuming each ***μ_k_*** known. We note that Johnson et al. (2008) also assumed a known core transition process. While we lose inference of uncertainty around core assignments and each ***μ_k_***, this problem has generally not been resolved in the literature and online estimation remains a major hurdle.

Finally, the multicore OU position likelihood is given by

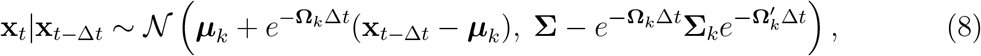

where **Ω**_*k*_ = *ω_k_***I**_2_ and 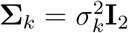 for the *k*th home range core.

#### Hierarchical inference across individuals

Full Bayesian inference about population level parameters can be obtained with the “Lunn method.” The Lunn method is a form of recursive Bayesian estimation and uses the marginal posteriors from a series of independent individual-level models fit with Markov chain Monte-Carlo (MCMC; or HMC) as the proposal distributions in a second stage MCMC algorithm (Lunn et al., 2013; Hooten et al., 2016; Hooten and Hefley, 2019). Here, to obtain population-level estimates of the population-level OU and core switching parameters, we can specify:

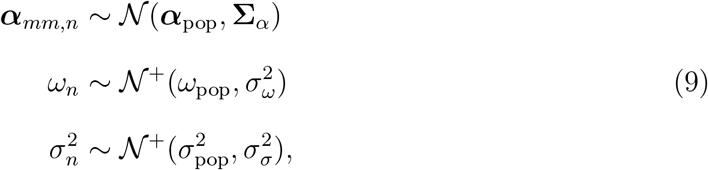

where ***α***_*mm,n*_ is the vector of coefficients correlating the core-switching covariates with staying in the *n*th individual’s most used core *m*, and **Σ**_*α*_ is a diagonal matrix of the among-individual variances for each covariate. It is convenient to restrict inference about ***α***_pop_ to the most-used core because individuals can have different numbers of core areas. Normal priors on each element of ***α***_pop_, truncated normal priors on *ω*_pop_ and *σ*_pop_, and inverse gamma priors on all among-individual (random effect) variances are conjugate priors and permit Gibbs updates for all population-level parameters. The individual-level parameters still require Metropolis-Hastings (MH) updates within the second stage algorithm, but these are straightforward because the MH ratios do not depend on the data (i.e. the data models cancel in the ratio; Lunn et al., 2013; Hooten et al., 2016; Hooten and Hefley, 2019).

### Application

#### Model system

Golden eagles are a long-lived, territorial raptor that reach sexual maturity entering their third breeding season (Kochert et al., 2002; Watson, 2010). They most commonly nest on cliffs, or less commonly large trees, and are generally central place foragers (Kochert et al., 2002; Watson, 2010). Eagles with established territories where a nest is a central place surrounded by uniformly average landscape should be expected to range and use space in a circular pattern around the nest. Because real landscapes are not uniform, an eagle’s realized space use would then be shaped by the habitat surrounding that central point. Primary prey of golden eagles nesting in Alaska are snowshoe hare *Lepus americanus*, ptarmigan *Lagopus* spp., and Arctic ground squirrel *Urocitellus parryii* (McIntyre and Adams, 1999; McIntyre and Schmidt, 2012; Herzog et al., 2019).

When a pair of eagles initiate a nesting attempt, the male does the majority of the provisioning, while the female tends the nest and does most of the incubating and brooding of eggs/nestlings. When nestlings mature to the point that they can thermoregulate (∼ 3 wk post-hatch; or when a nest fails), the adult female no longer needs to tend them as regularly, so she is free to move about the territory and aid in provisioning (Watson, 2010). We expect that this event should be commensurate with an abrupt change in space use, because nest-tending requirements suddenly become less restrictive. This might allow space use to change so that the male and female of the breeding pair partition space to minimize overlap in foraging areas and/or territory defense efforts. It is also possible that this might occur dynamically throughout the season and/or day, regardless of nest tending duties.

Another key characteristic of golden eagles that would be expected to strongly influence how they use space is their flight mechanics—they are a soaring bird capable of capturing dynamic air currents to decrease or completely offset the energetic costs of flight (Katzner et al., 2012; Watson, 2010). Consequently, their space use patterns, and possibly partitioning of space among individuals, will be shaped dynamically by weather variables (Eisaguirre et al., 2020). Two common forms of such flight subsidies are thermal uplift, caused by the sun heating the surface of the earth and causing air to rise, and orographic uplift, caused by wind blowing up slope.

Because habitat and weather features are non-uniform around nest sites/central places, eagles (and other animals) can establish multiple core areas within their larger home range. Thus real home ranges are not a single circular distribution in a homogeneous landscape, but multiple cores shaped by the non-uniform distribution of food and energy subsidies.

#### Telemetry data

We captured golden eagles with a remote-fired net launcher placed over carrion bait near Gunsight Mountain, Alaska (61.67^◦^N 147.35^◦^W). Captures occurred during spring migration, mid-March to mid-April 2016. Adult eagles were equipped with 45-g back pack solar-powered Argos/GPS platform transmitter terminals (PTTs; Microwave Telemetry, Inc., Columbia, MD, USA). PTTs were programmed to record a GPS location every other hour, yielding 12 fixes per day. Eagles were sexed molecularly and aged by plumage. See Eisaguirre et al. (2018) or Eisaguirre et al. (2019) for additional details.

#### Selection covariates

We used the Alaska Center for Conservation Science Alaska Vegetation and Wetland Composite (AKVWC; 30-m resolution) data for characterizing habitat type. We collapsed the numerous habitat types in the dataset into eight for this analysis. These were shrub, open (e.g., meadows and open tundra), bare, forest, wet (e.g., marsh), water, ice (i.e. perennial snow and ice), and human. See Appendix 1 for details. Elevation data were gathered using the Mapzen Terrain Service with the elevatr package (Hollister and Shah, 2018). We specified the ‘zoom’ variable such that the resolution closely matched that of the habitat data. We included elevation and slope (*slope* ∈ [0, *π/*2] radians) as predictors in the model.

We used a state-wide data set of snow-off date (date of which an area became snow free) to derive a dynamic binary indicator variable of whether or not grid cells were free of snow (Macander et al., 2015). While one might expect some confounding between the (perennial) snow and ice habitat variable and this snow indicator, it would be limited due to few glaciated and perennial snow-covered areas frequented by the eagles sampled. The remaining variables included in the model were related to orographic and thermal uplift and were derived from the National elevation data and Center for Environmental Predictions (NCEP) North American Regional Reanalysis (NARR) data. Angle of incidence (aoi) was included for the effect of orographic uplift on eagle space use. It is the deviation of the relative wind from the aspect of a slope and was computed such that *aoi* ∈ [0, *π*] (Murgatroyd et al., 2018); *π/*2 corresponds to a wind orthogonal to a slope’s aspect, and *π* to a wind perfectly parallel to a slope’s aspect thus blowing directly up slope. Wind direction was computed trigonometrically from the meridional and zonal wind components estimated by the NCEP NARR 10 m above the surface.

The effect of thermal uplift was included with a hill shade variable. Hill shade was computed following Murgatroyd et al. (2018), such that *hs* ∈ [0, 1], where *hs* = 1 is direct sun (most thermal uplift) and *hs* = 0 no sun (no thermal uplift). We gathered the required location-, date-, and time-specific azimuth and zenith of the sun using the package maptools (Bivand and Lewin-Koh, 2016).

#### Core switching covariates

We also included wind variables as covariates in the core transition process. We expected that certain wind directions and/or magnitudes might make certain home range cores more or less favorable. So, the cosine and sine of wind direction were included in addition to wind magnitude as covariates in equation (7). As above, these were computed trigonometrically from the NCEP NARR data specific to each home range core. Among-core distance was also included as a covariate to account for more frequent transitions to closer cores.

#### Estimation and inference

To illustrate our approach, we used only data from eagles that were clearly defending territories in 2016. This included six males and six females, all aged to their fifth year or older. None of these eagles were members of the same breeding pair. Aerial surveys flown in June 2016 revealed that four of the eagles had young (at the time of the survey), and, with the exception of one nest site that was not surveyed, the others showed signs of reproductive attempts.

Individual-level marginal posteriors of the core switching and OU process parameters were obtained using Stan (Stan Development Team, 2018). We used three chains of 5,000 iterations, including 2,000 for warmup, retaining every third sample for a total of 3,000 samples. These were then used as proposed values for the MH updates in the second stage for estimating parameters in equation (9). The population-level selection coefficients **β** were estimated with the empirical Bayes procedure with a Bayesian hierarchical logistic regression model in Stan (default normal priors; Stan Development Team, 2016), using marginal posterior predictive samples as available points, as in the simulations above. Convergence to the posterior was checked with trace plots and Gelman diagnostics (Stan Development Team, 2018). Stan and R code for fitting the individual-level OU process and sampling from the conditional posterior predictive distributions, as well as R code for the second stage MCMC algorithm, are provided in Appendix 2.

As our primary interest was in differences between male and female eagles in early and late breeding season, we wanted parameter estimates specific to each sex and to early and late breeding season. To keep computing time more reasonable, we fit the model separately and in parallel (on multiple CPU cores) for these periods as well as for each sex. Aerial observations of the nests of the tagged eagles indicated that 20 June was on average the approximate date when nestlings should have been of age to thermoregulate, so we used this date to partition the data between early and late breeding season.

Utilization distributions were computed according to equation (6). The probability density predicted for each home range core was weighted by the number of eagle locations in that core prior to computing the 95% volume contour of the space use distributions, which we used to estimate home range boundaries (Hooten et al., 2017).

## Results

### Movement parameters

Because individuals had differing numbers of home range cores, we present here only the OU movement parameters from the individuals’ most heavily used core. We found a slight increase in centralizing tendency for males (Fig. 3) and an increase in the number of home range cores for both sexes from early to late breeding season (Fig. S3). We found a weak effect of stronger wind correlating with females staying in their most-used (nesting) core and wind direction affecting males’ propensity to stay within that core during early breeding season (Fig. 3).

**Figure 3:**
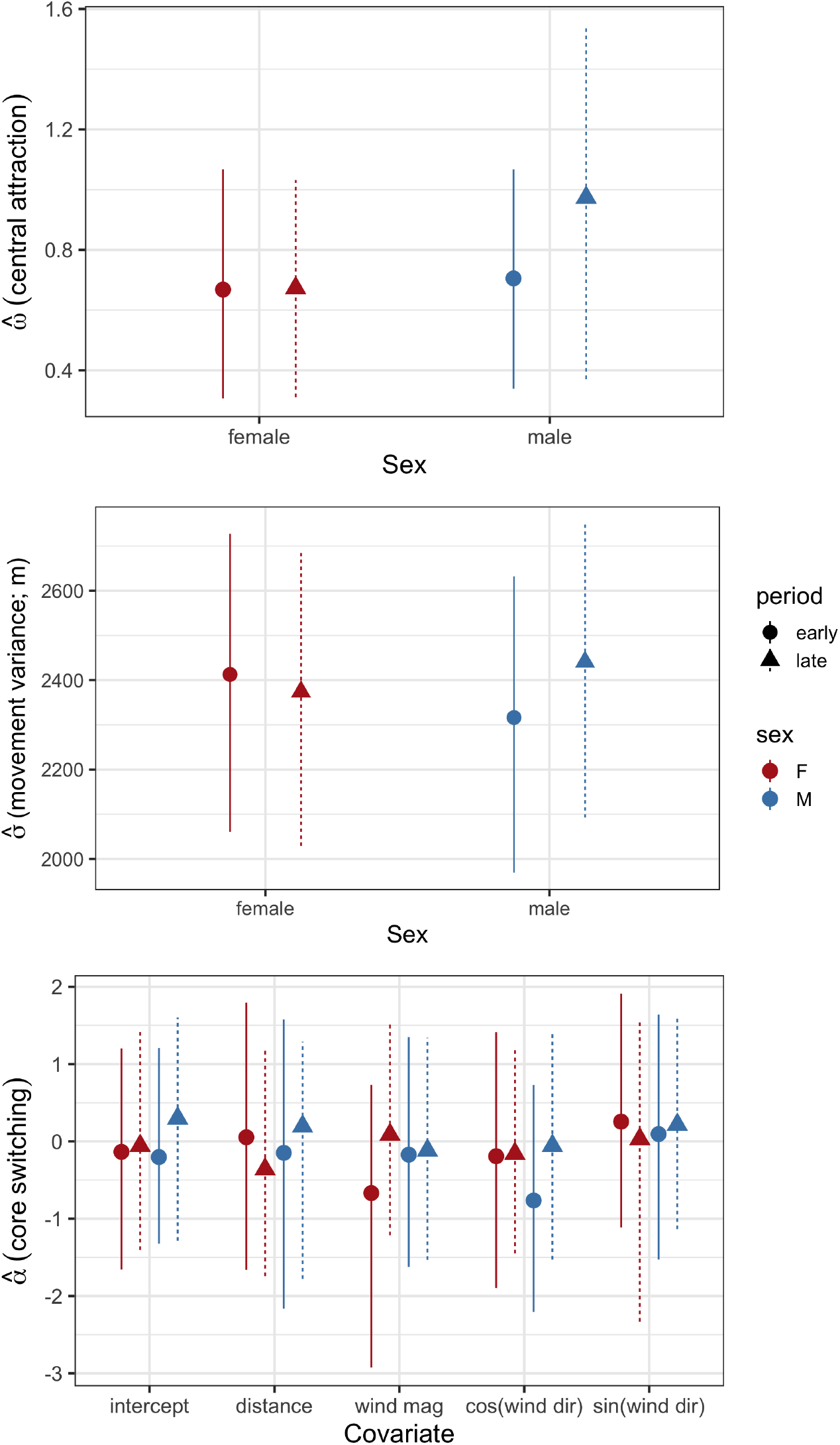
Posterier means and 90% credible intervals of the popoulation-level movement parameters in an Ornstein-Uhlenbeck movement model fit to six male and six female golden eagles with territories in southcentral Alaska. *σ* is the movement variance; *ω* the autocorrelation parameter measuring the centralizing tendency; and ***α*** the coefficients in the Markovian home range core switching process correlating the covariates to staying in the most used home range core. The models were fit separately for early and late breeding season.

### Habitat selection

We present the effects of the most relevant habitat types in figure 4, which comprised *>* 99% of the space used (see figure S5 in Appendix 1 for all habitat types). Both male and female eagles weakly selected against forested areas during early breeding season, and females selected against shrub and open habitats early, relative to bare areas (Fig. 4). Overall, males and females used similar terrain, though there was some evidence that females used slightly steeper slopes (Fig. S4).

**Figure 4:**
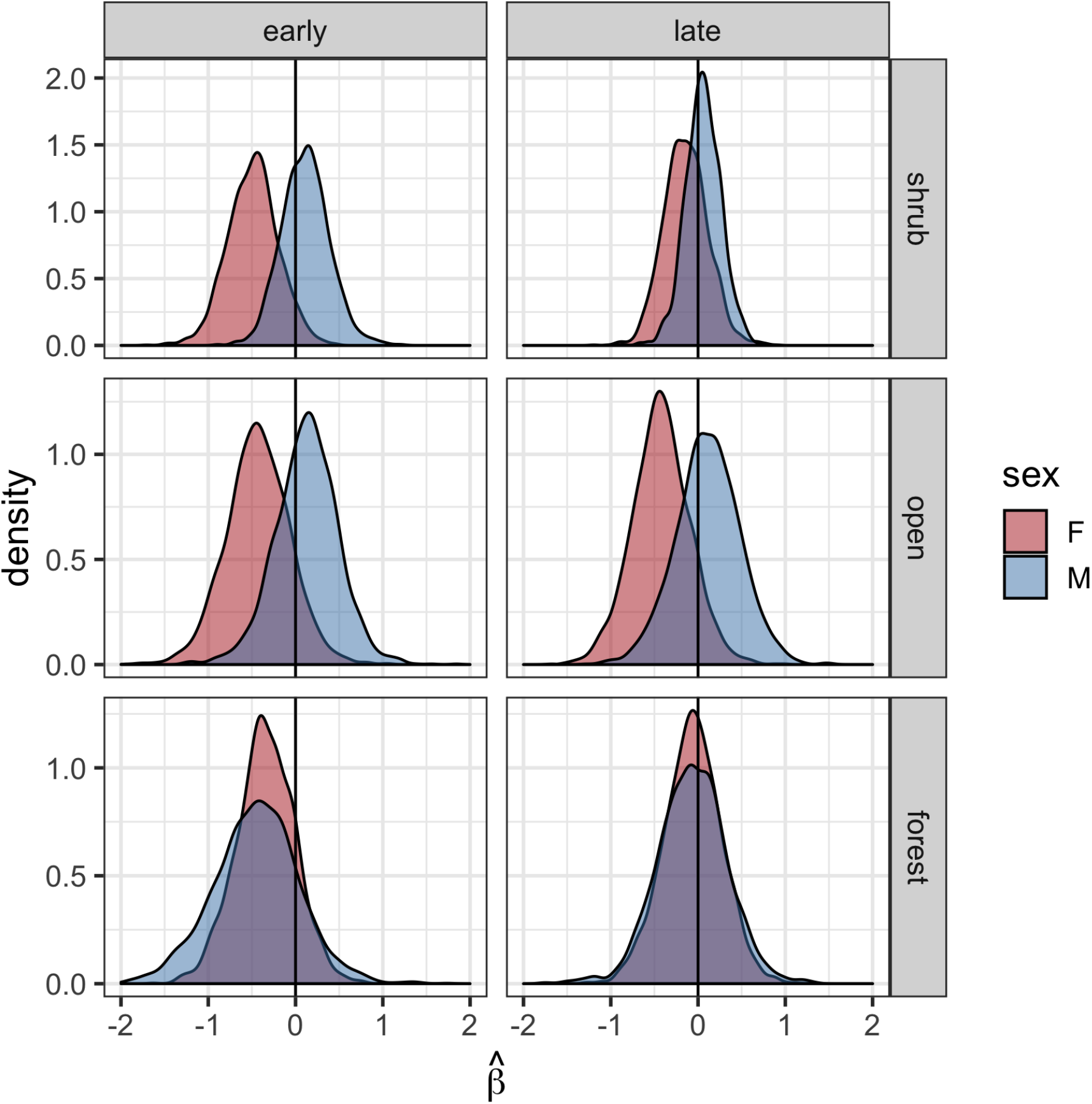
Marginal posterior densities of the population-level habitat selection parameters showing partitioning of certain habitat types by male and female golden eagles. These were estimated with an Ornstein-Uhlenbeck space use model for territorial golden eagles summering in southcentral Alaska. Densities were constructed with 2000 posterior samples. The reference category used for estimation was ‘bare’.

### Energy landscape

In early breeding season, before nestling thermoregulation or nest failure, males and females appeared to select energy landscape features similarly (Fig. 5–7). During late breeding season, male and female eagles appeared to partition the landscape dynamically based on components of the energy landscape (Fig. 5–7). Males tended to use areas with more orographic uplift (i.e. higher angle of incidence; Fig. 5), while females used more thermal uplift (i.e. greater hill shade; Fig. 5). This pattern resulted from males and females selecting dynamic energy subsidy features over the landscape differently (Fig. 6). Further, females showed essentially no selection for or against angle of incidence during late breeding season (Fig. 5 & 6). The posterior probability that females selected more strongly for hill shade than males was 0.94, and the posterior probability that males selected more strongly for higher angle of incidence than females was 0.82 (Fig. 6). These probabilities were computed relative to the posterior mean for the opposite sex with Monte Carlo integration.

**Figure 5:**
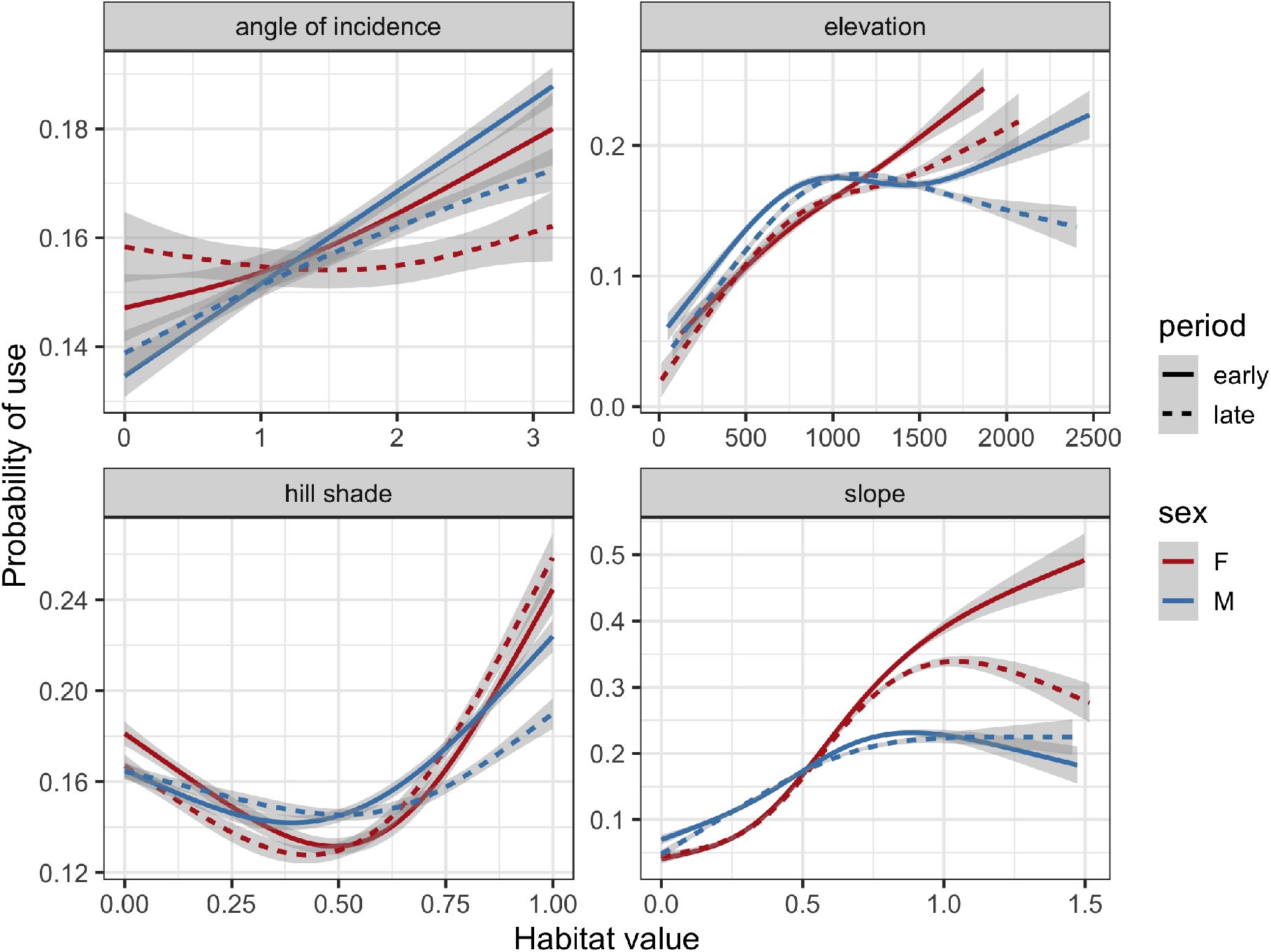
Probability of a golden eagle using a spatial location within its breeding season home range in southcentral Alaska as a function of habitat variables estimated with an Ornstein-Uhlenbeck (OU) space use model. This is the average effect conditioned on the space available to each eagle characterized by an OU biased random walk. The model was fit separately for early and late breeding season and for each sex. Predictions were smoothed over the availability points with a generalized additive model (*df* = 4) and ribbons are 95% confidence intervals. Units are radians for angle of incidence and slope, and meters for elevation. Higher hill shade corresponds to more direct sun and greater thermal uplift potential. We present a version of this figure with common *y*-axis scales in Appendix 1.

**Figure 6:**
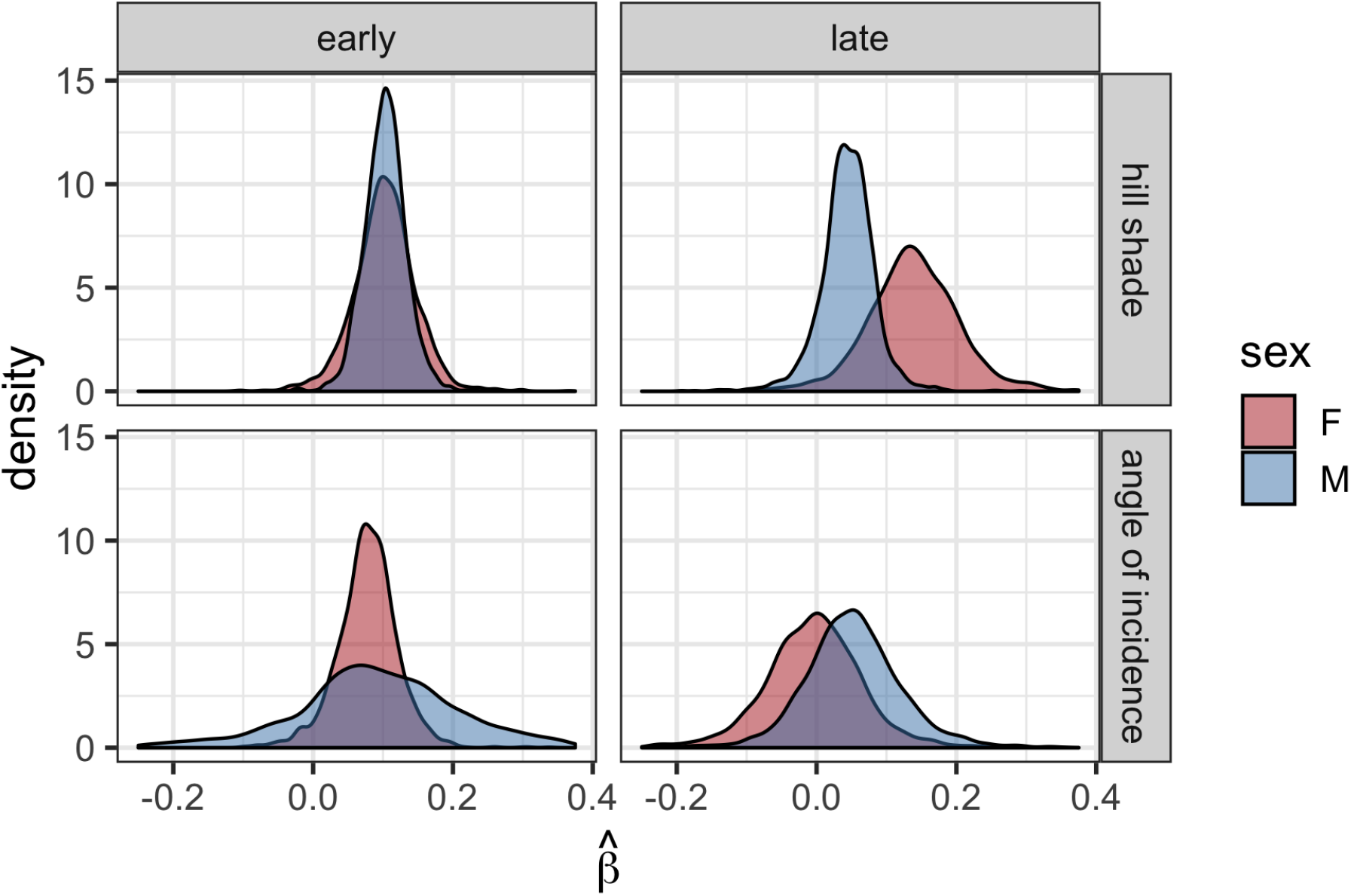
Marginal posterior densities of population-level hill shade and angle of incidence selection parameters showing partitioning of the energy landscape (thermal and orographic uplift) by male and female golden eagles during late breeding season. These were estimated with an Ornstein-Uhlenbeck space use model for territorial golden eagles summering in southcentral Alaska. Densities were constructed with 2000 posterior samples.

## Discussion

Here, we demonstrated a method that overcomes analytical and computational challenges in fitting a hierarchical mechanistic home range model to data but also, as we showed through a simulation study, provides unbiased inference about biologically interpretable parameters. The OU space use model allows inference about movement behavior, resource selection, and, ultimately, space use patterns, and it is applicable to any central place animal. Further, we demonstrated that this approach can be extended to account for additional complexity in the structure of animal home ranges—in the form of multiple core areas—and possible covariates affecting transitions within that structure. Applying the model to real data offered novel insight into the movement and space use of an organism that is sensitive to its central place, landscape resources, and energy subsidies available in a fluid atmosphere.

### Application to the energy home range

Applying the OU space use model to territorial golden eagle movement revealed some notable patterns. First, male and female eagles had relatively similar space use patterns during early breeding season, followed by a shift at the approximate time of a particular phenological event. When nestlings are able to thermoregulate, the female of a pair can take on additional duties (i.e. provisioning and territorial defense). Our results show this coincides with a change in space use, emergent from changes in both resource selection and movement behavior.

Male and female eagles partitioned their use of orographic and thermal uplift during late breeding season (Fig. 5 & 7). Two possible explanations for this are that it (1) serves as a means for each sex to avoid overlap in foraging and/or territory defense efforts and/or (2) is an emergent pattern resulting from size dimorphism. Reverse sexual size dimorphism is prevalent in raptors, and it is coupled with dimorphism in wing loading (i.e. wing area per body mass): Females of many raptors, including golden eagle, exhibit higher wing loading than males (Lish et al., 2016). Lighter wing loading could allow male eagles to capitalize on even slight bits of uplift generated orographically with more energetic efficiency than females. Thermal uplift is also generally a more efficient flight subsidy than orographic uplift (Duerr et al., 2012), so, given their higher wing loading, it is likely energetically advantageous for females to use primarily thermal soaring. Similar patterns have been recently shown in sexually size dimorphic wandering albatrosses *Diomedea exulans*, where males—the sex with higher wing loading—favor flight in more energetically favorable wind conditions than females (Clay et al., 2020).

**Figure 7:**
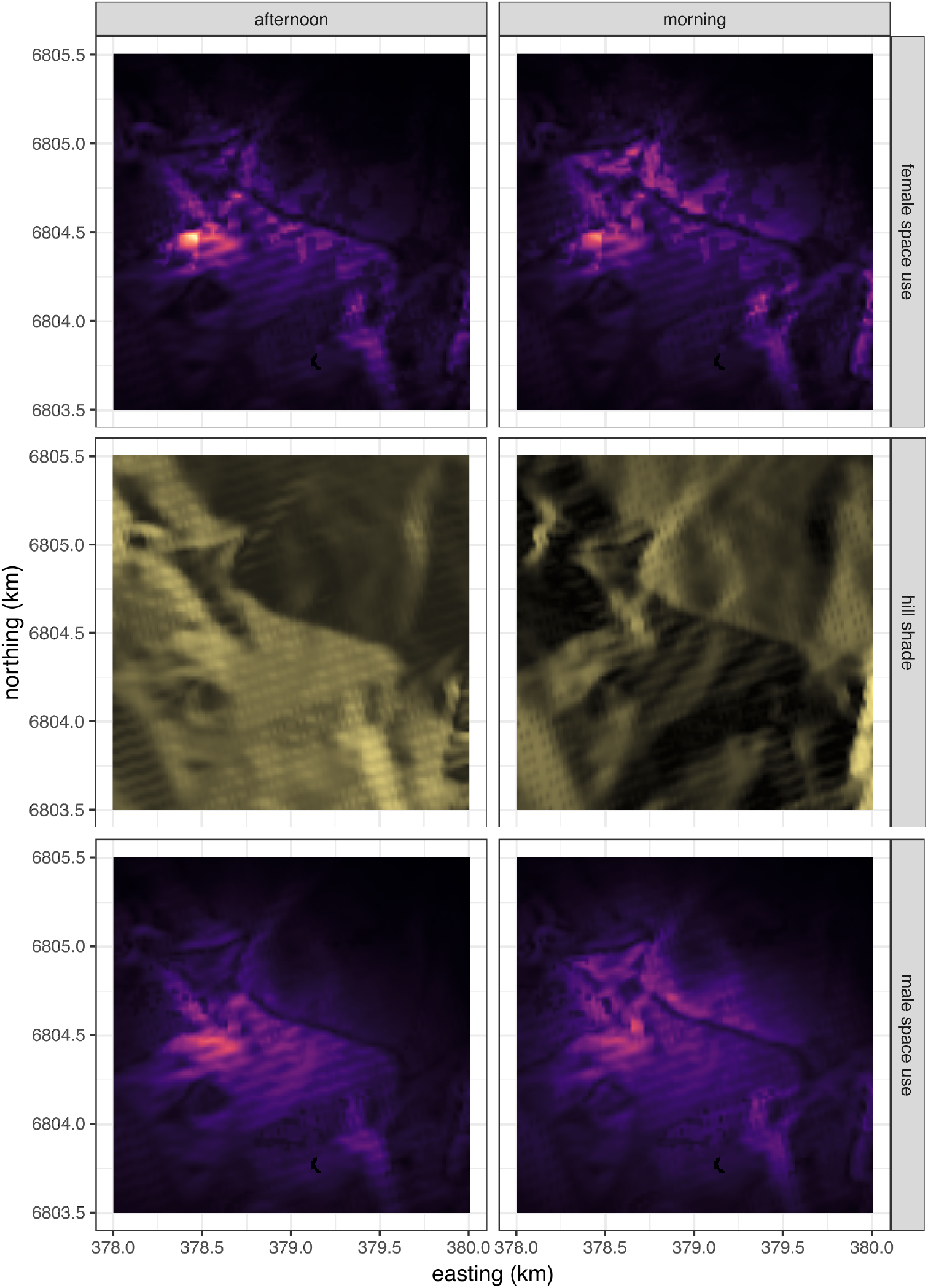
Hill shade maps and utilization distributions *f_u_*(**z**_*t*_*) predicted from the Ornstein-Uhlenbeck space use model for territorial golden eagles summering in south-central Alaska. Predictions were made over a characteristic landscape **z**_*t*_* during morning and afternoon to illustrate differential space use patterns according to thermal uplift. White corresponds to highest probability of use and black lowest.

Our results also suggest that soaring birds could dynamically segregate space vertically, as well as partition activity budgets. Orographic uplift is typically available at only relatively low heights above Earth’s surface, whereas thermals can travel much higher into the atmospheric boundary layer. The altitude of eagles using these different types of uplift follows suit (Katzner et al., 2015). Given selection for differing types of uplift, we would thus expect male and female eagles might also partition their home ranges vertically. Maintaining good visibility with the surface is required for successful foraging, so partitioning thermal and orographic uplift could also indicate different behavioral budgets. Further, thermal and orographic uplift vary over space following changes in wind and sun angle. Consequently, males and females may partition three dimensional space and activities temporally through the day, as females await better thermal soaring conditions before beginning extensive movements around the home range. In contrast, wind can generate orographic uplift at any time during the day.

While our findings relating to the energy landscape were most notable, we also found some differences in habitat and terrain use, which are consistent with sex-specific roles during the breeding season. Females used and selected steeper slopes than males, consistent with nesting behavior and perching near the nest (Collopy and Edwards, 1989; Kochert et al., 2002; Watson, 2010). Not surprisingly, females used less steep slopes during late breeding season, compared to early, consistent with behavior in the later nestling stages of breeding (Collopy and Edwards, 1989; Watson, 2010). Also, males, who do most of the provisioning even late into the breeding season (Collopy and Edwards, 1989; Watson, 2010), selected more strongly for shrub and open habitats (Fig. 4), which would likely be used for hunting. During late breeding season, females’ selection for shrub habitats approached that of bare areas, likely following an increased role in provisioning.

### Modeling central place space use

Our modelling approach is conceptually framed around the elastic disc hypothesis, an analogy underlying central place theory, and it shares and integrates aspects of a number of other methods. It is analytically similar to the general frameworks presented by Johnson et al. (2008) and Christ et al. (2008); however, it overcomes estimation difficulties by implementing the model analogously to common RSF and SSF approaches (Manly et al., 2002; Forester et al., 2009; Avgar et al., 2016; Hooten et al., 2017). Computationally, it is far simpler to implement but produces similar parameter estimates and biological inference. Additionally we extended the model to the cases where covariates may drive the use of multiple home range core areas and population-level inference across multiple individuals (i.e. partial pooling) is needed. Further, we demonstrated how recursive Bayesian estimation can be particularly useful in estimating complex, computationally demanding hierarchical movement models (Lunn et al., 2013; Hooten et al., 2016; Hooten and Hefley, 2019; Hooten et al., 2019).

The OU space use model, as we and others have shown, yields unbiased inference about resource selection parameters (Johnson et al., 2008). This is despite inherent identifiability issues in studying the movement of central place animals. That is, it is difficult to identify whether an animal uses its central place disproportionately to other space because (1) it must tend the central place, (2) there is favorable habitat there, or (3) some combination of both. Unfortunately, this can bias some of the movement parameter estimates (Fig. S1 & S2 Johnson et al., 2008); however, evidence suggests this occurs in only select cases (i.e. when spatial autocorrelation and selection are very high; Fig. S1 & S2; Johnson et al., 2008). Bias in 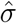 is potentially problematic because it additionally biases estimation of home range size. However, we found that this occurs when **β** and spatial autocorrelation are particularly high (Fig. S2)—higher than what we estimated in our application to real data (Fig. 4 & 6).

A tempting way of accounting for central places within conventional RSF and SSF approaches is to simply include ‘distance to central place’ in *ψ*(**z**). However, this assumes the central place is some feature of the landscape **z**, which is problematic for two reasons. First, it is inconsistent with central place behavior. The central place fundamentally modifies the animal’s behavior and space use; the animal does not actively select the central place as a resource while moving within its home range. Second, the selection coefficient *β* weighting this distance covariate would be biased high, as the central place is inappropriately discounted in *f_a_*(**z**), leaving *β* as the only parameter to make up for the disproportionate space use at the central point. In contrast, the OU space use model incorporates the central place such that it is an element on the landscape that modifies *f_a_*(**z**)—the animal’s movement and behavior.

The OU process can be presented as an advection-diffusion equation, as in equation (1); however, its properties are somewhat unique (Blackwell, 1997). Various other forms of advection-diffusion equations can also be used to mechanistically model home ranges and territories with central place dynamics (Moorcroft and Lewis, 2006), and these have been formally reconciled with resource selection analyses (Moorcroft and Barnett, 2008). Due to equation (6), however, when the OU process is integrated into the weighted distribution framework, steady state space use (or utilization) distributions are straightforward to compute. If we were to use an advection-diffusion process other than an OU process (e.g., different biased random walk or correlated random walk), computationally-intensive numerical investigation of the so-called master equation or simulations would be required (Moorcroft and Lewis, 2006; Barnett and Moorcroft, 2008; Potts et al., 2012, 2014a,b,c; Potts and Lewis, 2014; Signer et al., 2019). With our approach, it is therefore much more straightforward to visualize the effects of dynamic resources on space use over hypothetical and/or real landscapes (Fig. 7).

## Conclusions

Here we show that estimating a hierarchical mechanistic space use model is relatively flexible and can be eased by breaking into stages without sacrificing inference. While our approach is not without shortcomings—primarily discounting uncertainty in some components—we believe it is a step towards practical implementation of more complex movement and resource selection models that can improve our understanding of animal movement ecology. Our application provides evidence of dynamic sex-specific partitioning of the energy landscape within home ranges, as well as movement and habitat selection patterns consistent with eagle biology. While the model works most naturally with central place animals, the ability to incorporate multiple home range cores and the range of movement and space use patterns that can be captured with the OU parameters make it broadly applicable.

## Acknowledgements

T. & D. Hawkins, M. Kohan, B. Robinson, and many others provided support in the field, and J. Liguori and N. Paprocki helped age eagles. R. Barry, J. McIntyre, M. Short, and L. Berman provided helpful comments on a draft of the manuscript. To these friends, we are most grateful. Additionally, we thank two anonymous reviewers who helped greatly improve the manuscript. Funding was provided by the Alaska Department of Fish & Game (ADF&G) through the federal State Wildlife Grant Program. JME was supported by the Calvin J. Lensink Fund during part of the project.

## Data accessibility

All movement data used for this manuscript are archived in the online repository Move-bank (https://www.movebank.org/; ID 17680093). The data contain information considered confidential and sensitive by the State of Alaska (State Statute 16.05.815(d)), but they could be made available for research at the discretion of the Alaska Department of Fish & Game.

## Appendix 1 supplementary tables and figures

**Figure S1:**
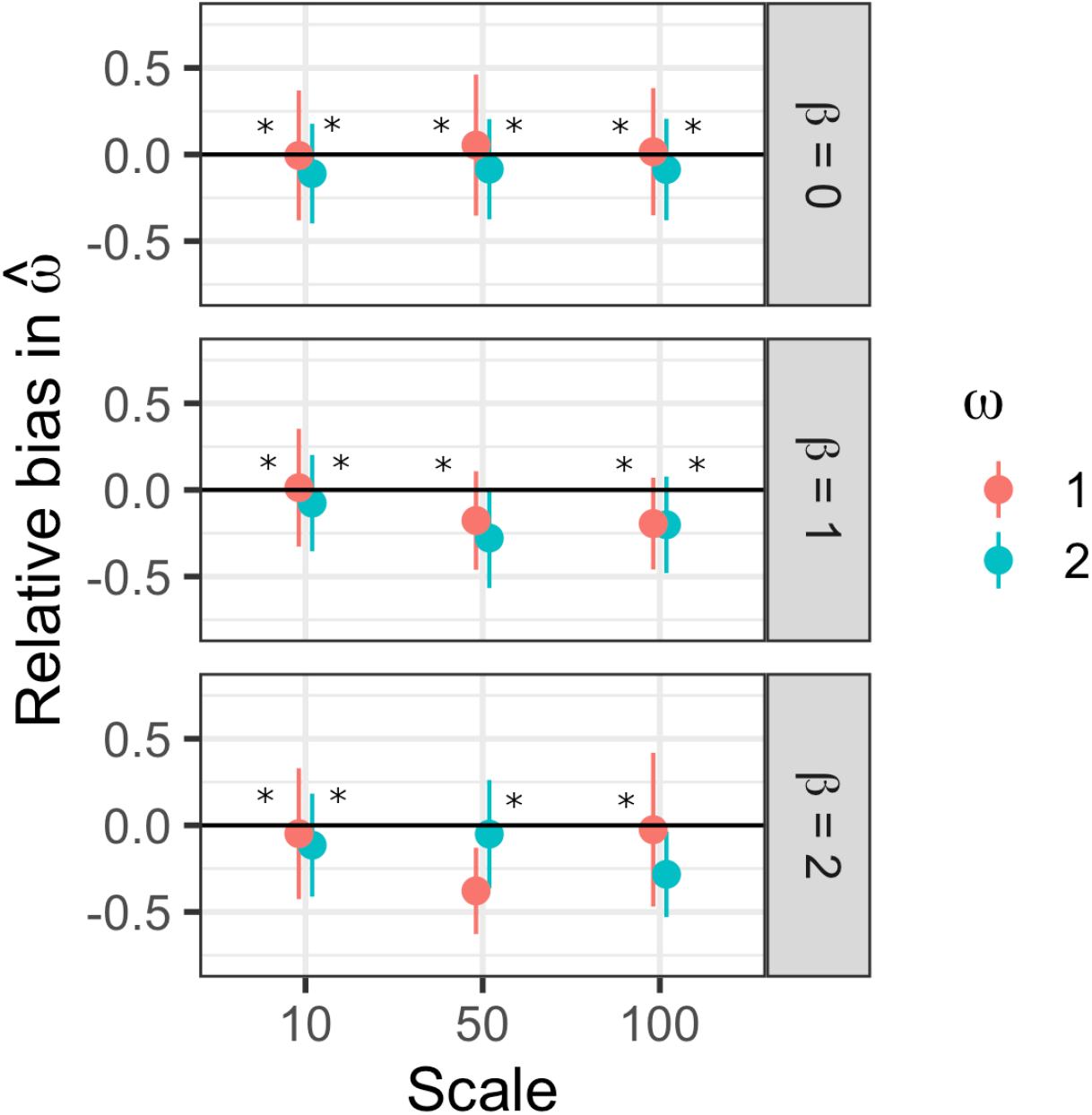
Relative bias in centralizing tendency when estimated with Ornstein-Uhlenbeck home range model with movement parameters estimated offline. Asterisk indicates 95% credible set captured the true value in *>* 70% of the simulations.

**Figure S2:**
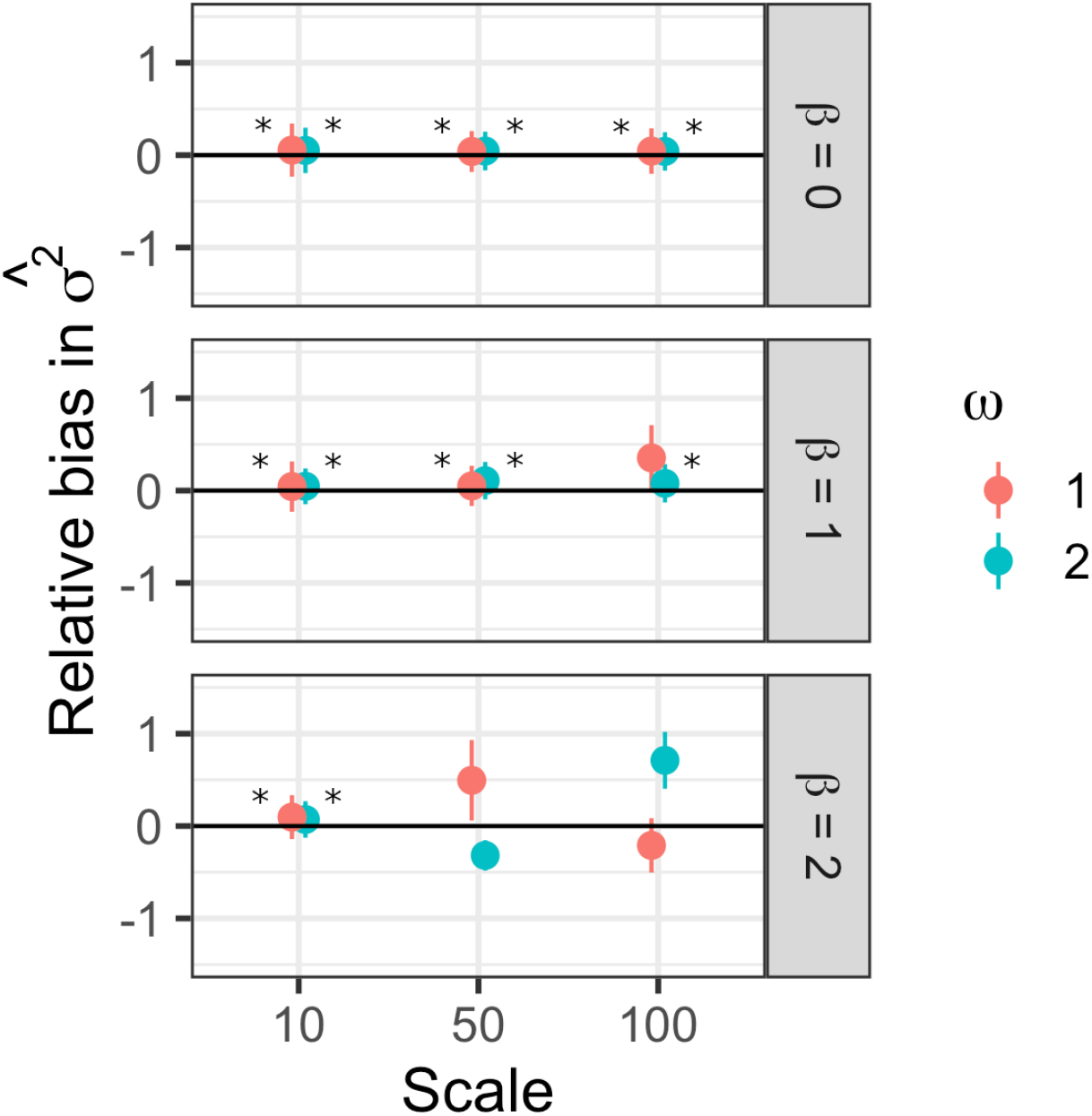
Relative bias in movement variance when estimated with Ornstein-Uhlenbeck home range model with movement parameters estimated offline. Asterisk indicates 95% credible set captured the true value in *>* 70% of the simulations.

**Table S1:**
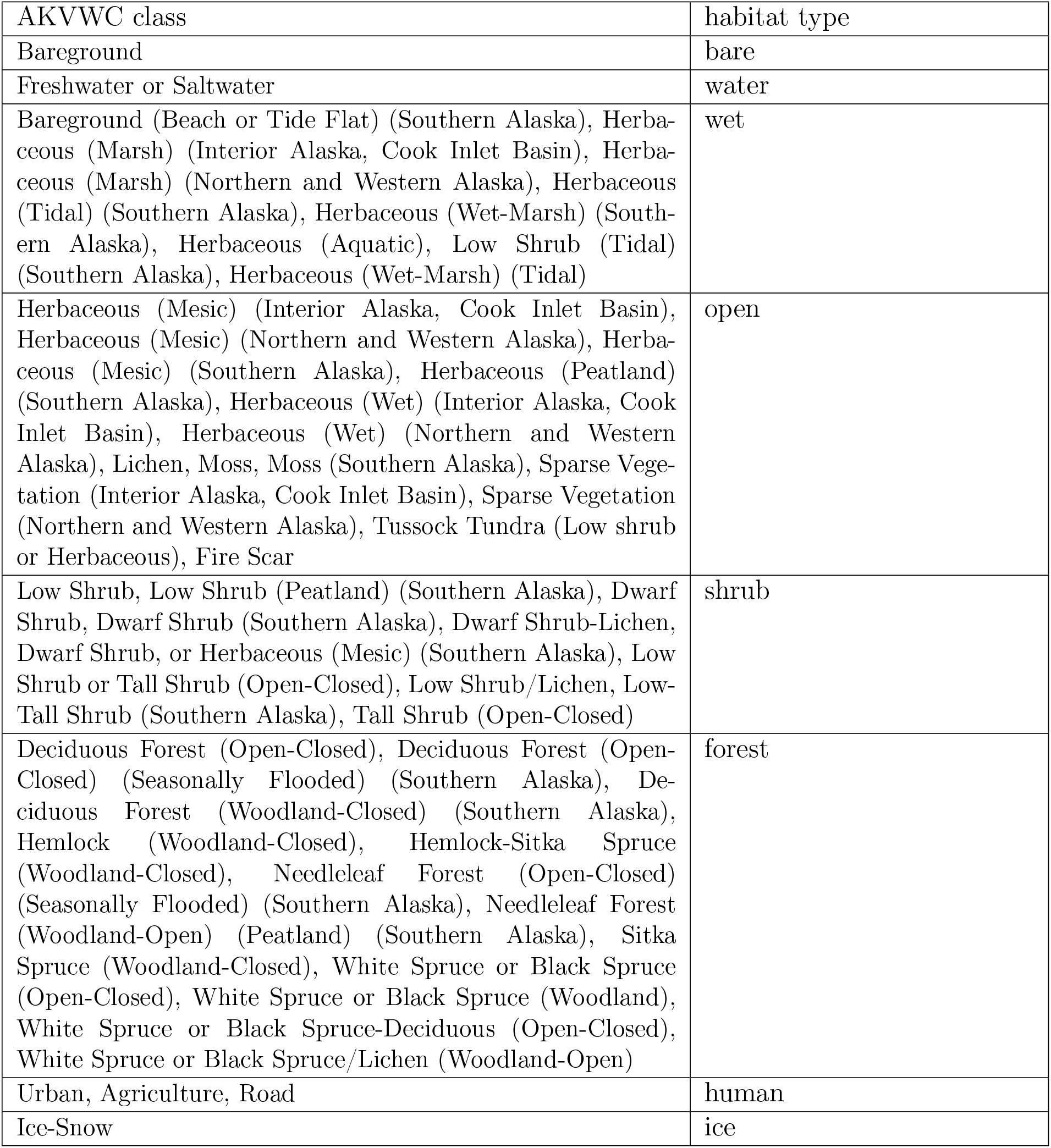
Habitat types used in analysis.

| AKVWC class | habitat type |
| --- | --- |
| Bareground | bare |
| Freshwater or Saltwater | water |
| Bareground (Beach or Tide Flat) (Southern Alaska), Herbaceous (Marsh) (Interior Alaska, Cook Inlet Basin), Herbaceous (Marsh) (Northern and Western Alaska), Herbaceous (Tidal) (Southern Alaska), Herbaceous (Wet-Marsh) (Southern Alaska), Herbaceous (Aquatic), Low Shrub (Tidal) (Southern Alaska), Herbaceous (Wet-Marsh) (Tidal) | wet |
| Herbaceous (Mesic) (Interior Alaska, Cook Inlet Basin), Herbaceous (Mesic) (Northern and Western Alaska), Herbaceous (Mesic) (Southern Alaska), Herbaceous (Peatland) (Southern Alaska), Herbaceous (Wet) (Interior Alaska, Cook Inlet Basin), Herbaceous (Wet) (Northern and Western Alaska), Lichen, Moss, Moss (Southern Alaska), Sparse Vegetation (Interior Alaska, Cook Inlet Basin), Sparse Vegetation (Northern and Western Alaska), Tussock Tundra (Low shrub or Herbaceous), Fire Scar | open |
| Low Shrub, Low Shrub (Peatland) (Southern Alaska), Dwarf Shrub, Dwarf Shrub (Southern Alaska), Dwarf Shrub-Lichen, Dwarf Shrub, or Herbaceous (Mesic) (Southern Alaska), Low Shrub or Tall Shrub (Open-Closed), Low Shrub/Lichen, Low-Tall Shrub (Southern Alaska), Tall Shrub (Open-Closed) | shrub |
| Deciduous Forest (Open-Closed), Deciduous Forest (Open-Closed) (Seasonally Flooded) (Southern Alaska), Deciduous Forest (Woodland-Closed) (Southern Alaska), Hemlock (Woodland-Closed), Hemlock-Sitka Spruce (Woodland-Closed), Needleleaf Forest (Open-Closed) (Seasonally Flooded) (Southern Alaska), Needleleaf Forest (Woodland-Open) (Peatland) (Southern Alaska), Sitka Spruce (Woodland-Closed), White Spruce or Black Spruce (Open-Closed), White Spruce or Black Spruce (Woodland), White Spruce or Black Spruce-Deciduous (Open-Closed), White Spruce or Black Spruce/Lichen (Woodland-Open) | forest |
| Urban, Agriculture, Road | human |
| Ice-Snow | ice |

**Figure S3:**
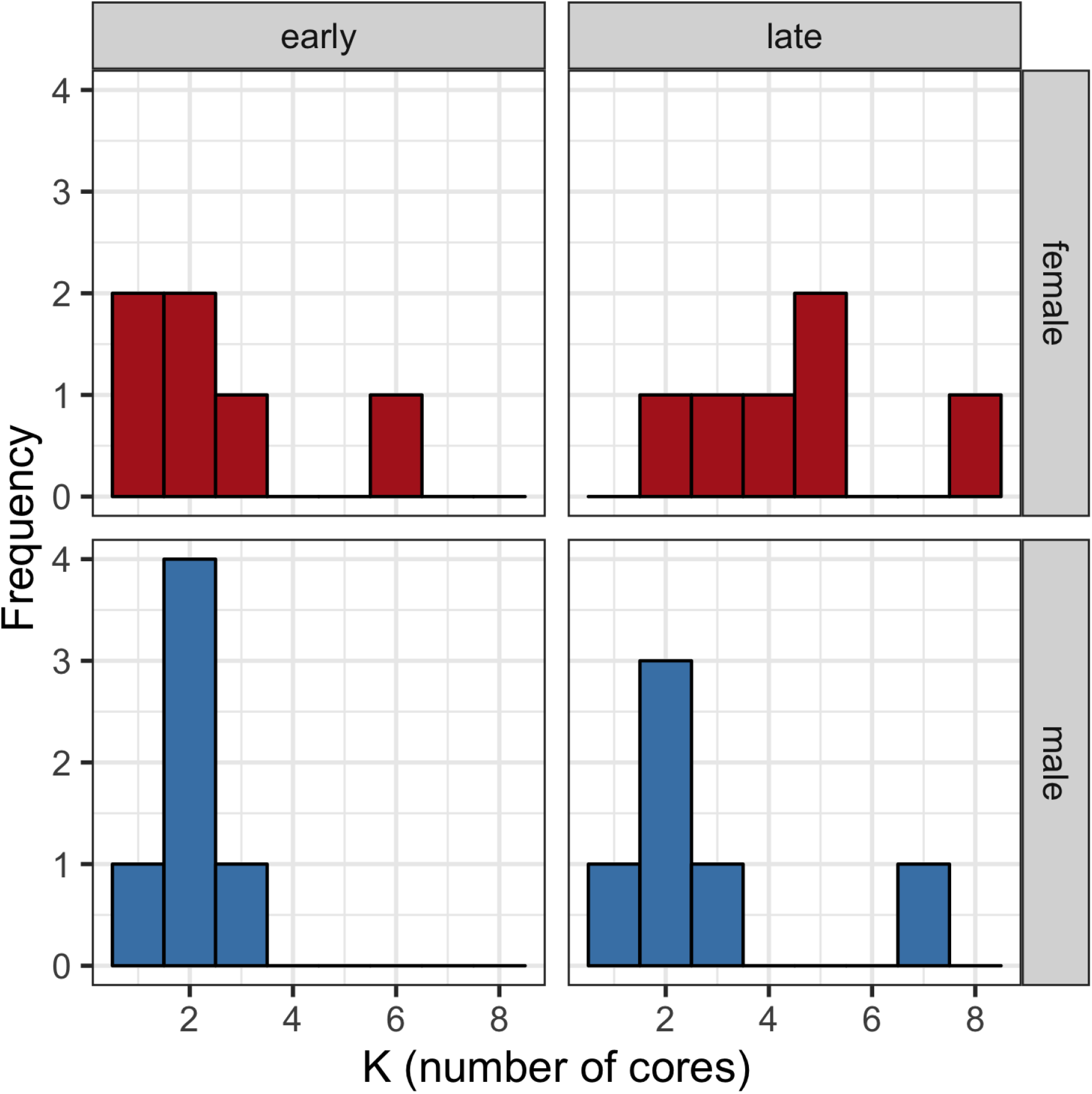
Number of home range cores estimated with a *k*-means clustering algorithm for six male and six female golden eagles with territories in southcentral Alaska. The algortithm was run separately for early and late breeding season.

**Figure S4:**
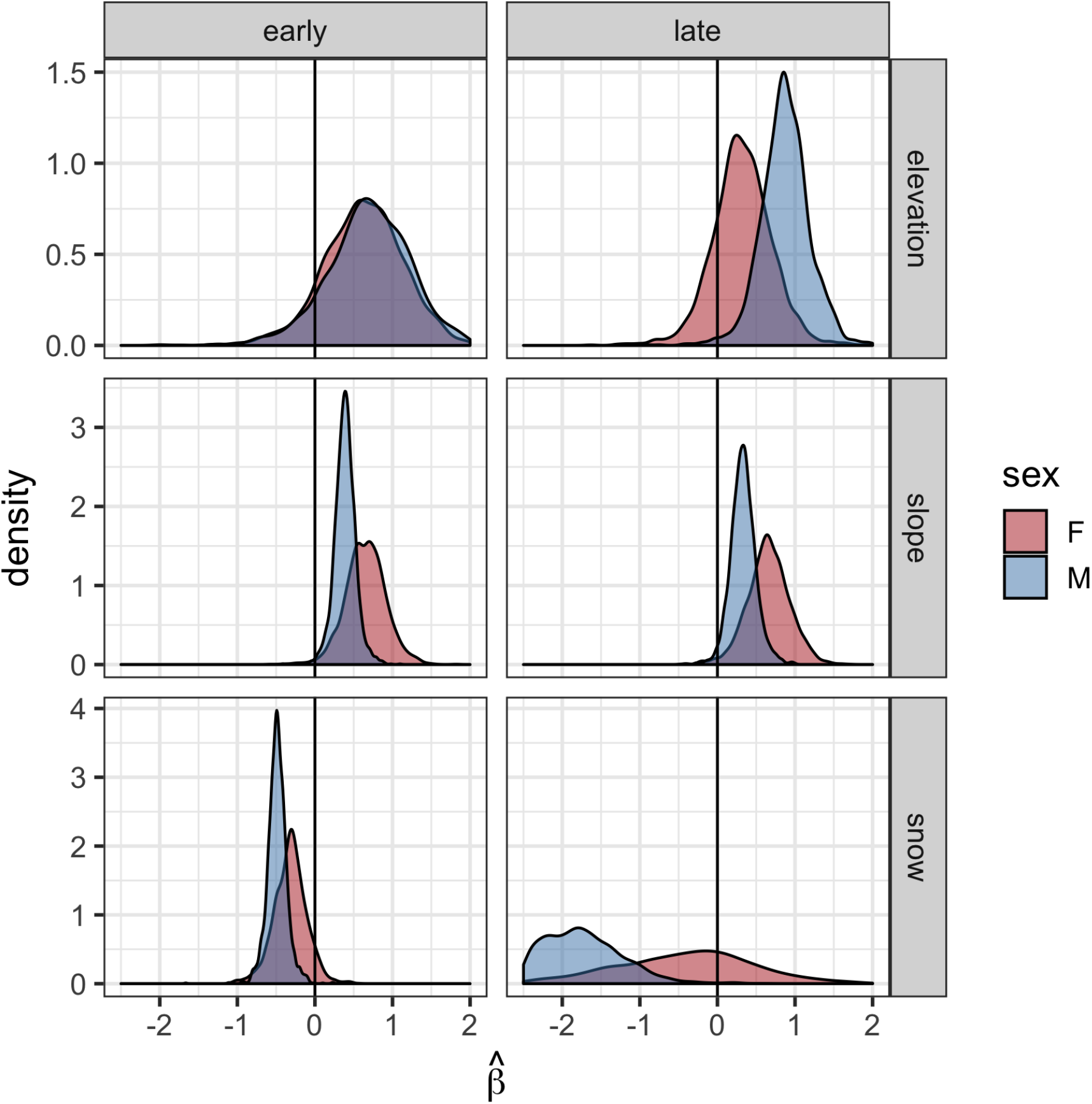
Marginal posterior densities of the population-level habitat selection parameters showing partitioning of certain habitat types by male and female golden eagles. These were estimated with an Ornstein-Uhlenbeck space use model for territorial golden eagles summering in southcentral Alaska. Densities were constructed with 2000 posterior samples. The snow variable was a dynamic indicator of whether or not a location was snow-free. The reference category used for estimation was ‘bare’.

**Figure S5:**
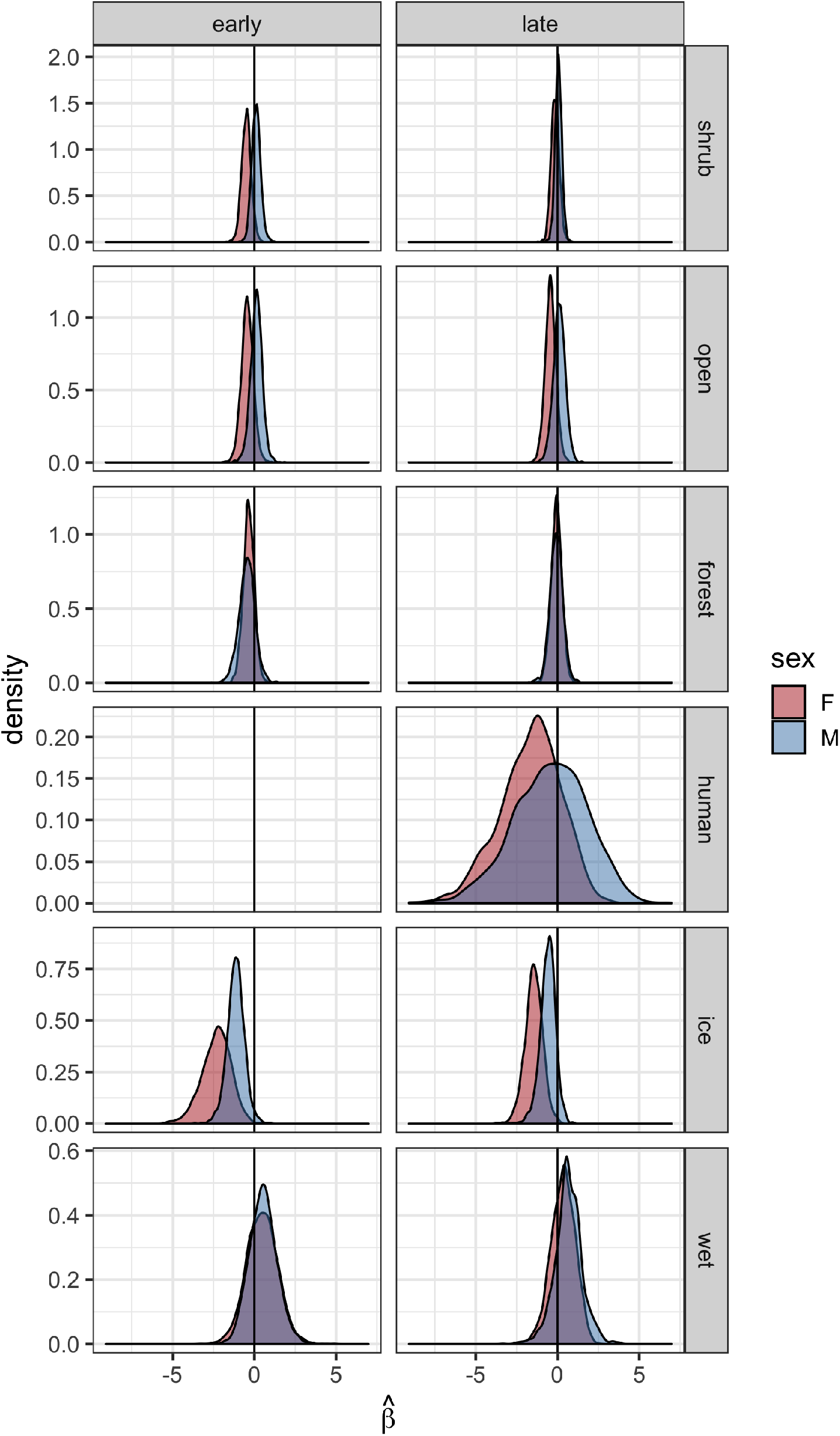
Full version of figure 4 from main text.

**Figure S6:**
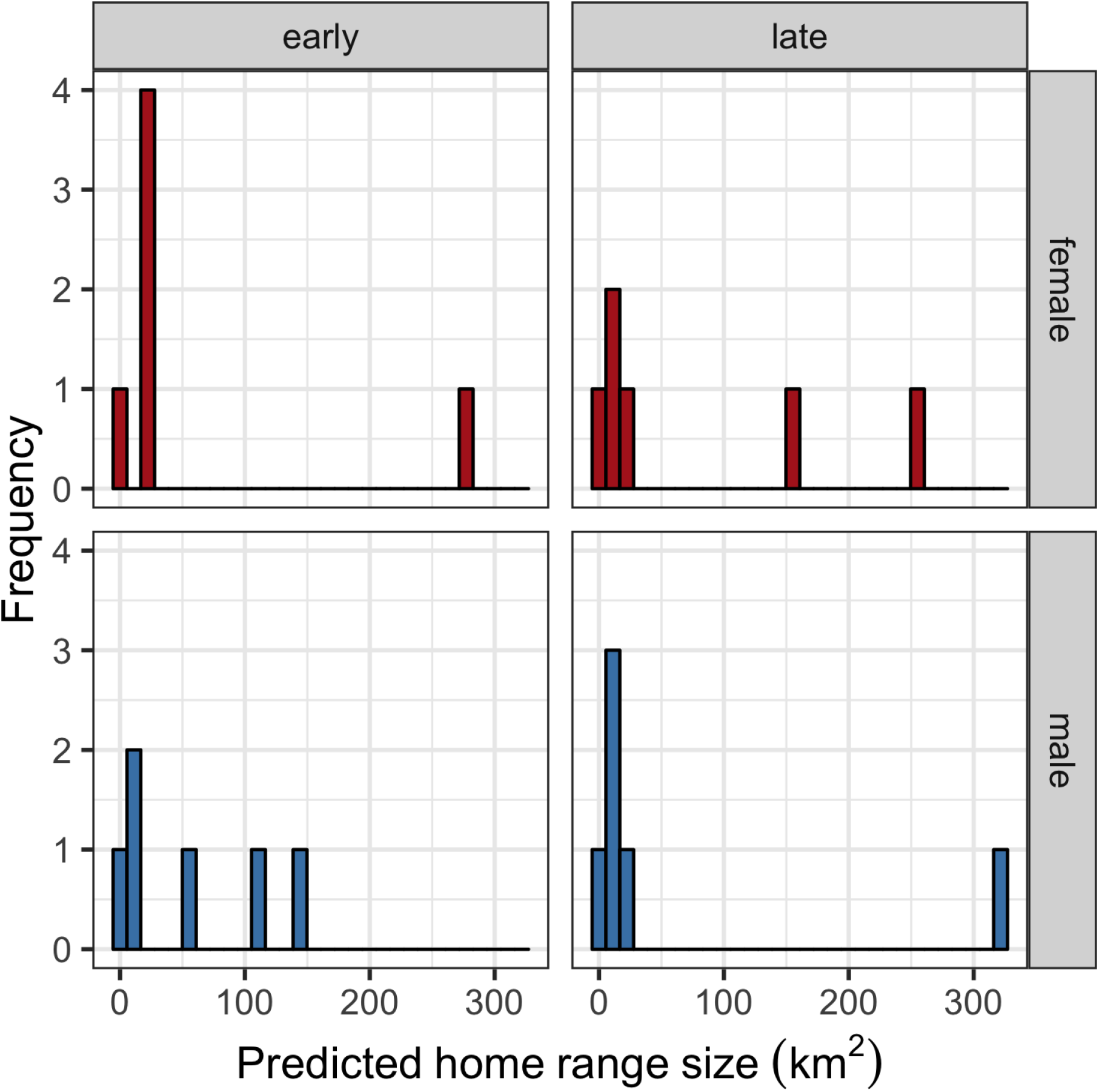
Home range sizes predicted from the Ornstein-Uhlenbeck space use model for territorial golden eagles summering in southcentral Alaska. Home range size was estimated as the 95% volume contour of the predicted space use distribution.

**Figure S7:**
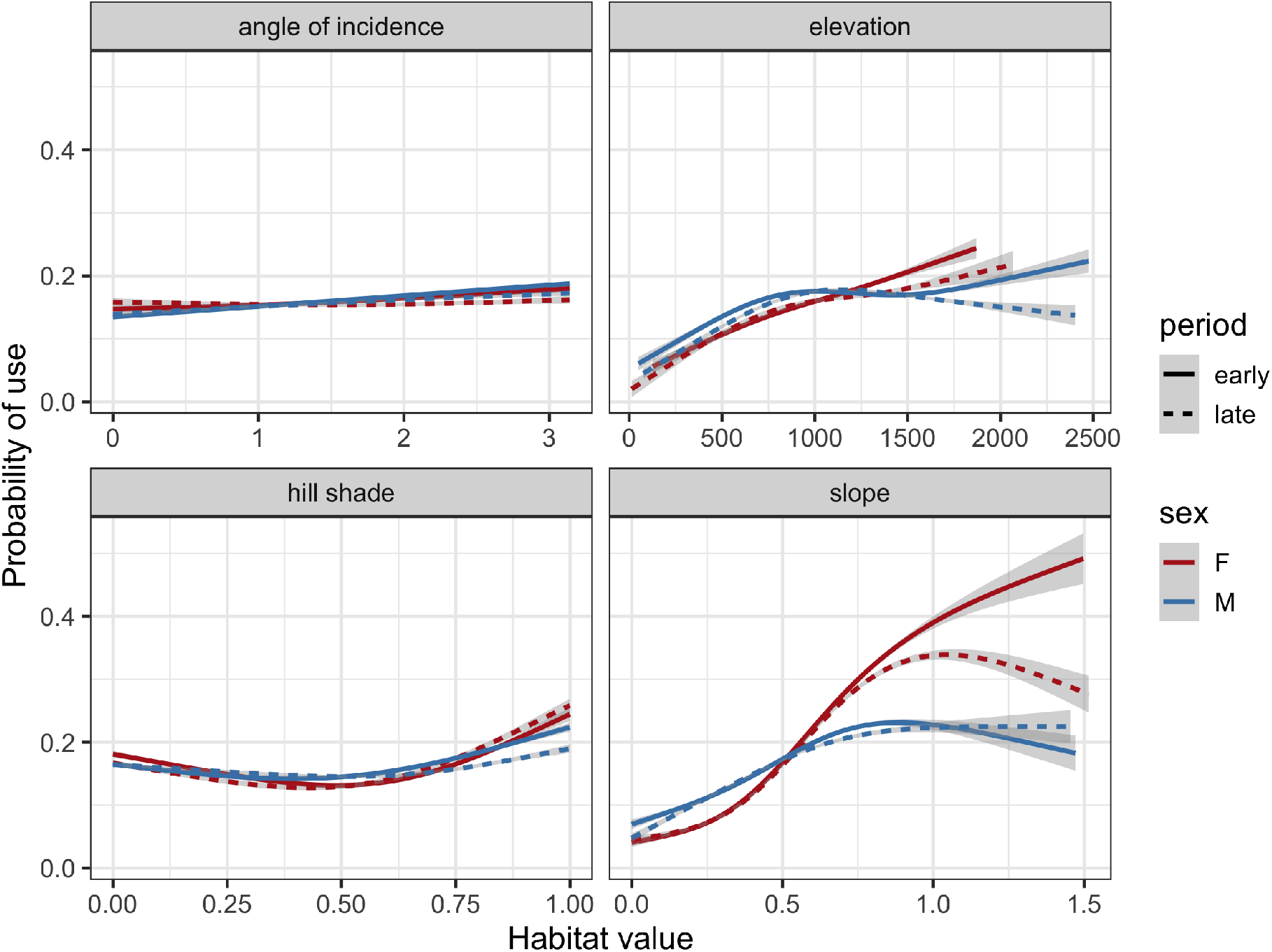
Version of figure 5 from the main text with common *y*-axis scales.

**Figure S8:**
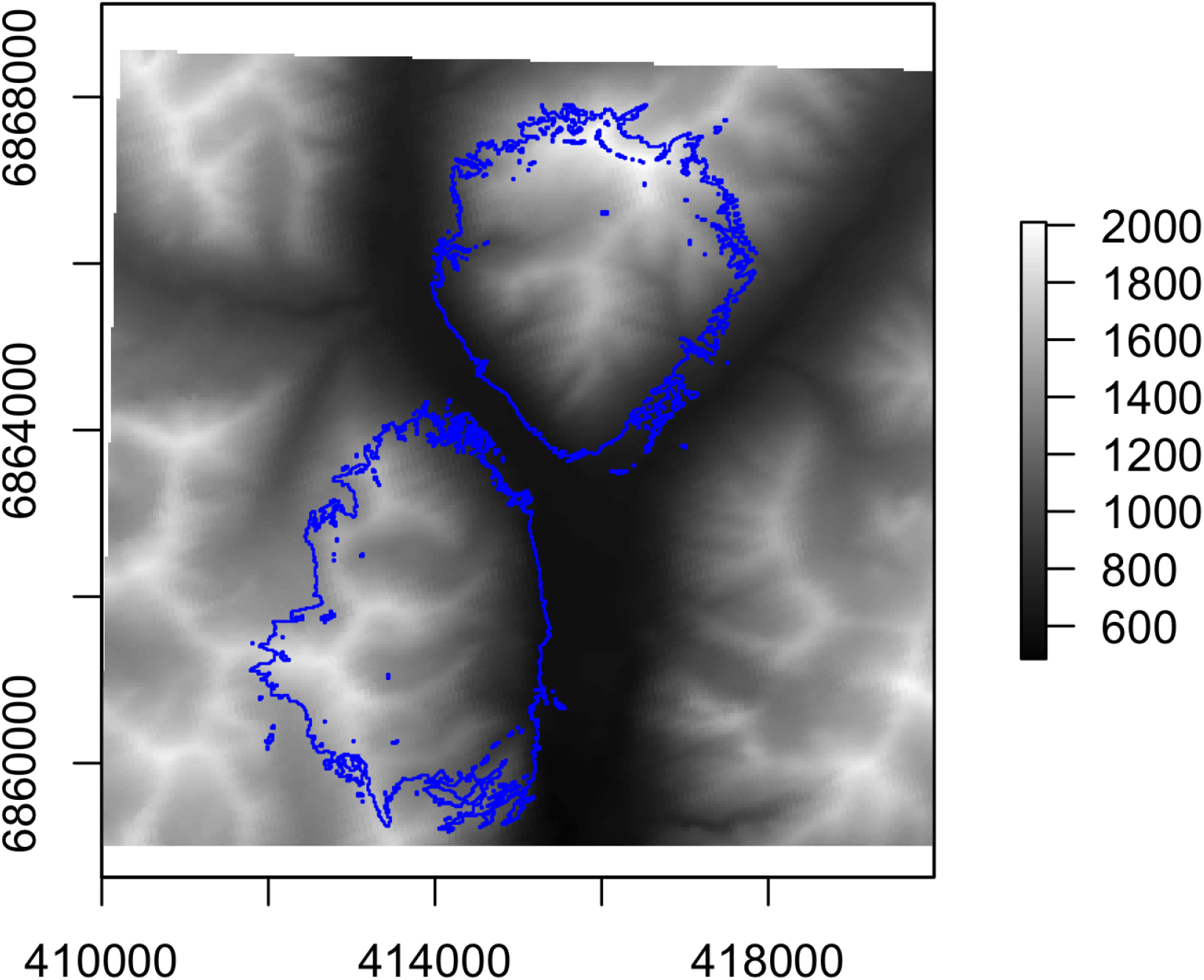
Example of predicted home range with two core areas. Home range boundary is the 95% volume contour of the predicted utilization distribution from the OU space use model. Base map is elevation. All units are meters.

## Appendix 2 code

## Stan model

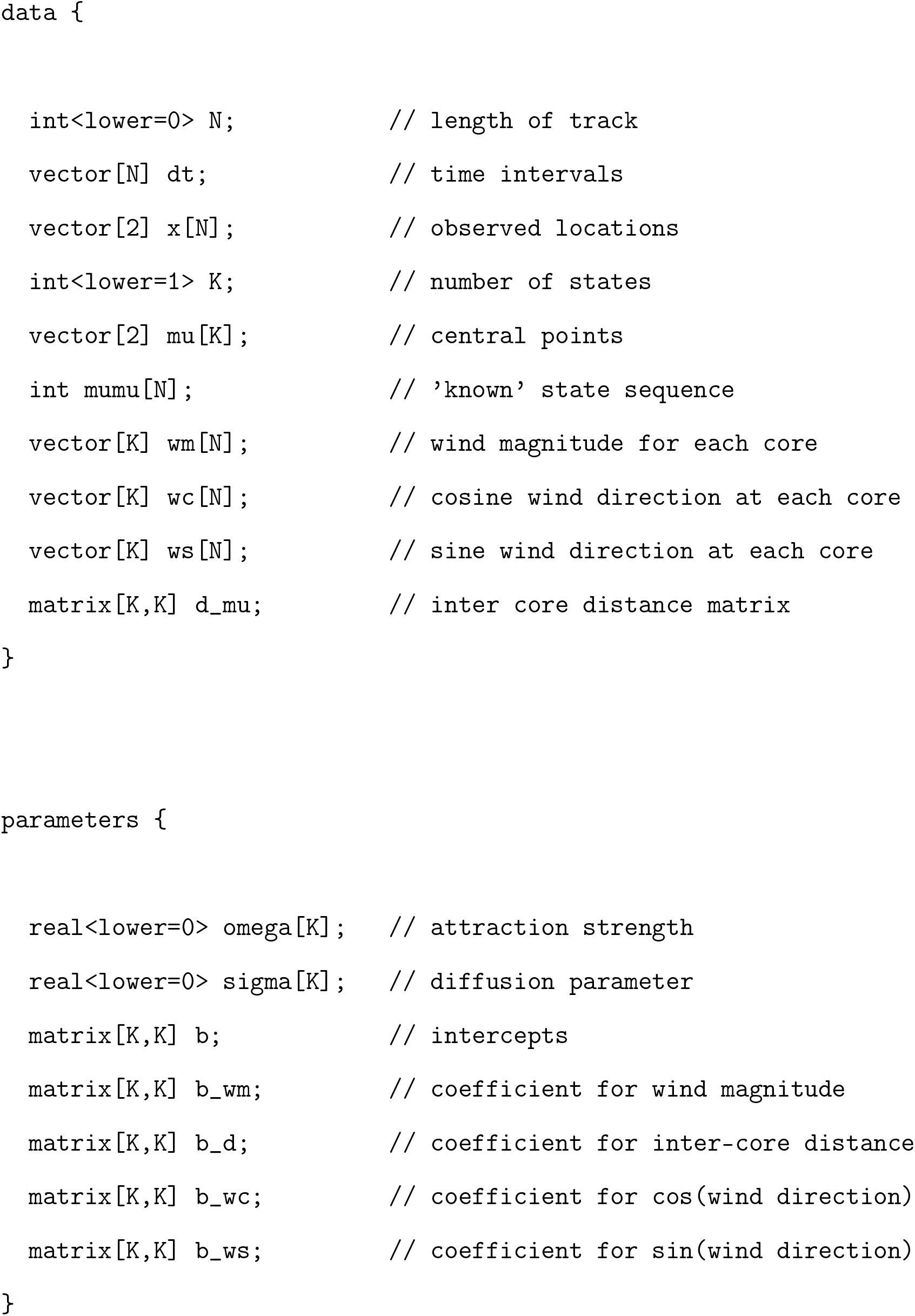
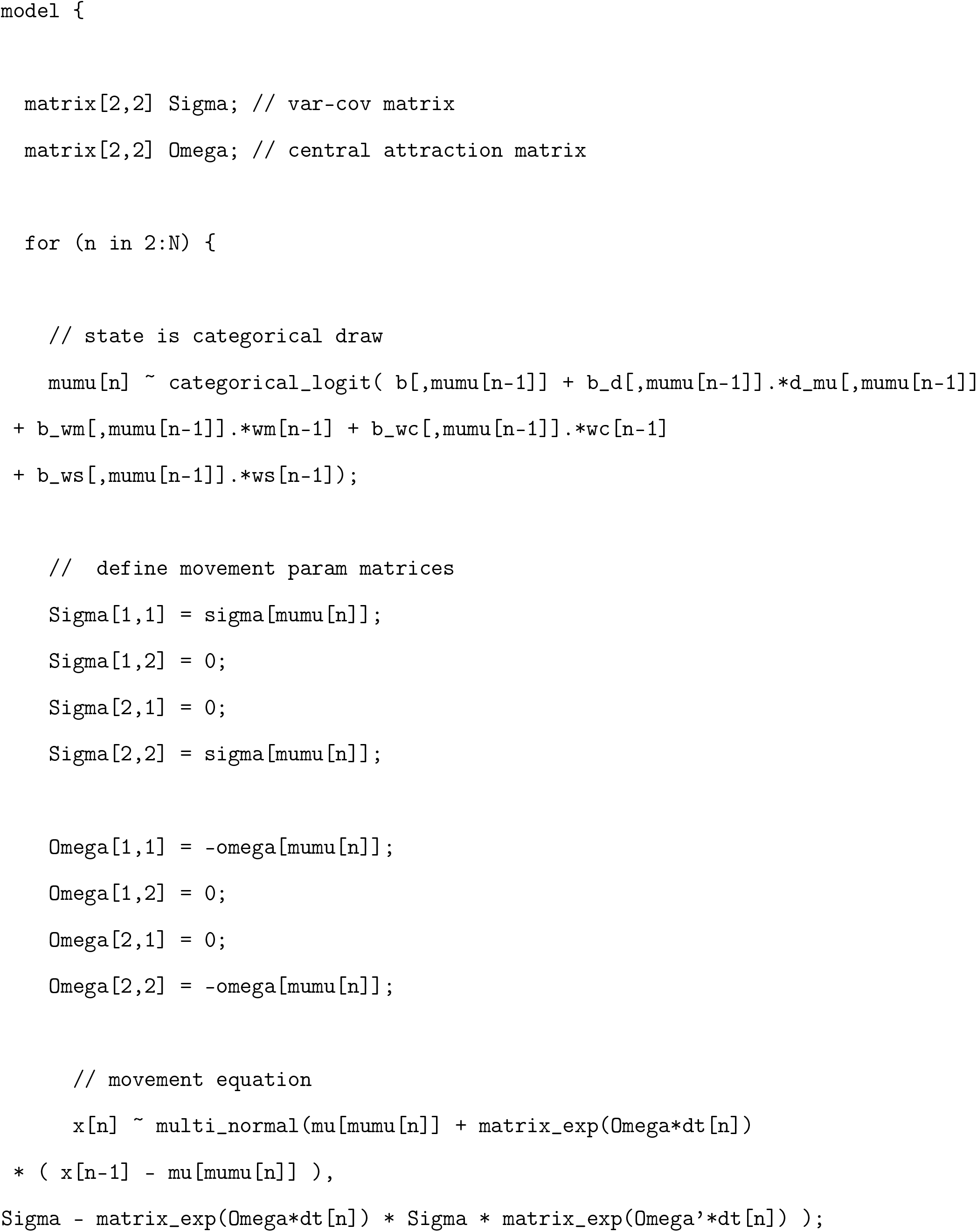
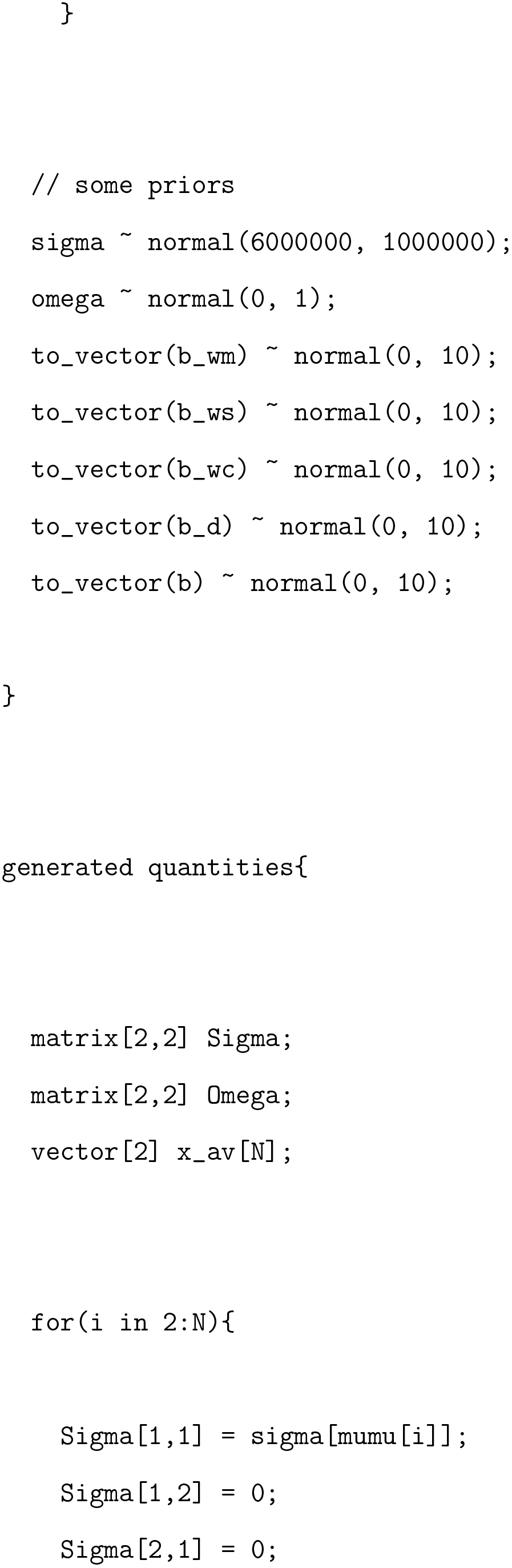
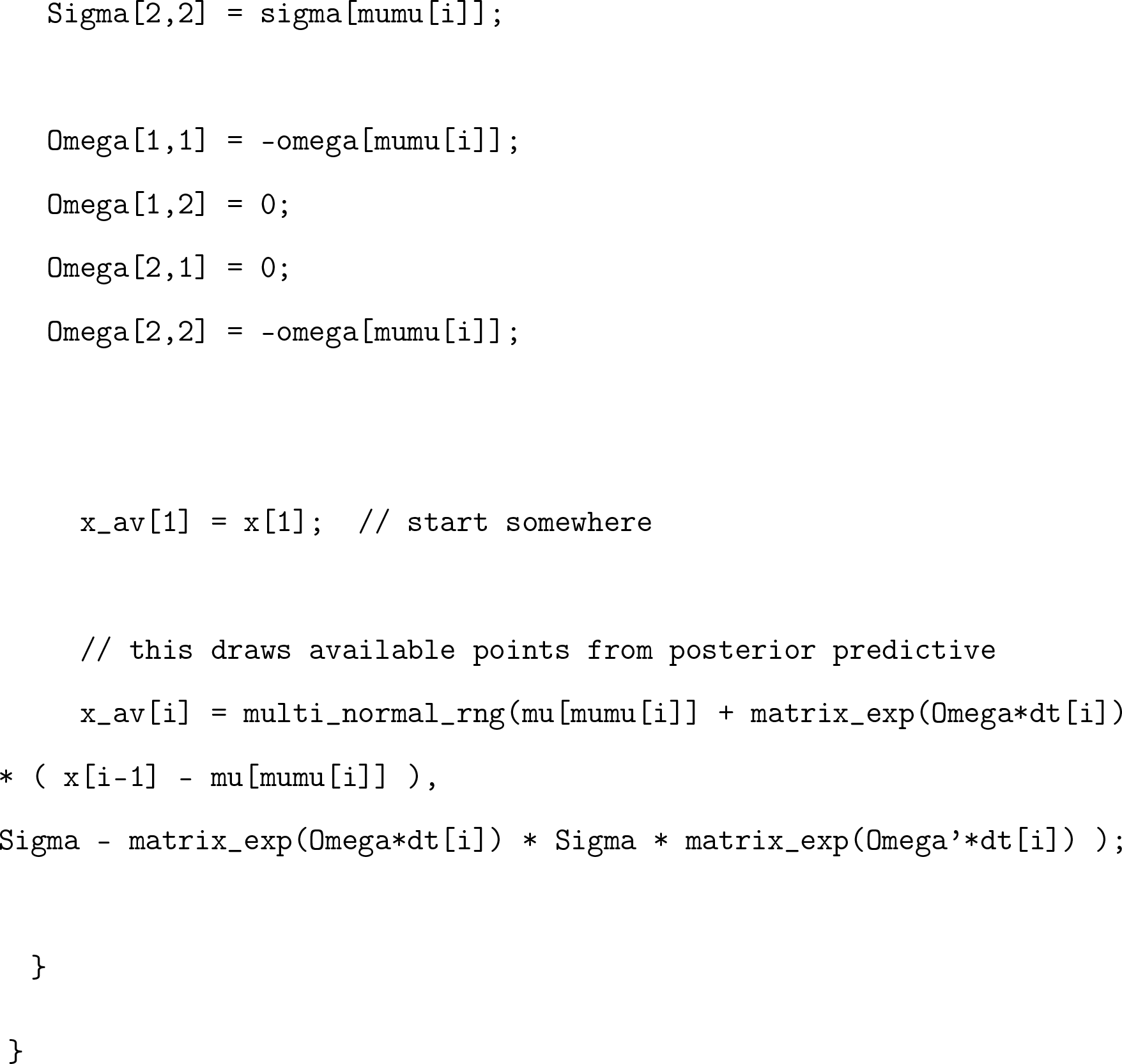

## R code

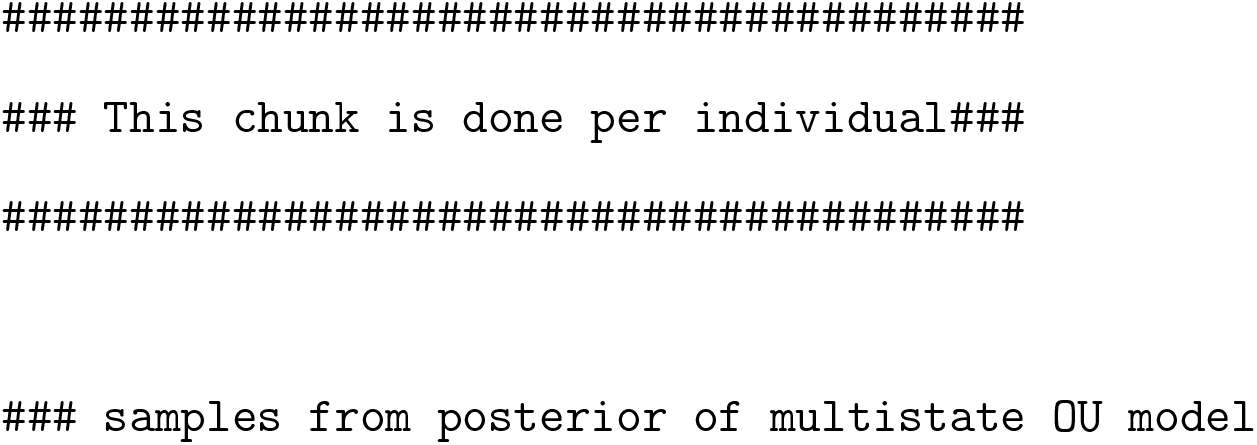
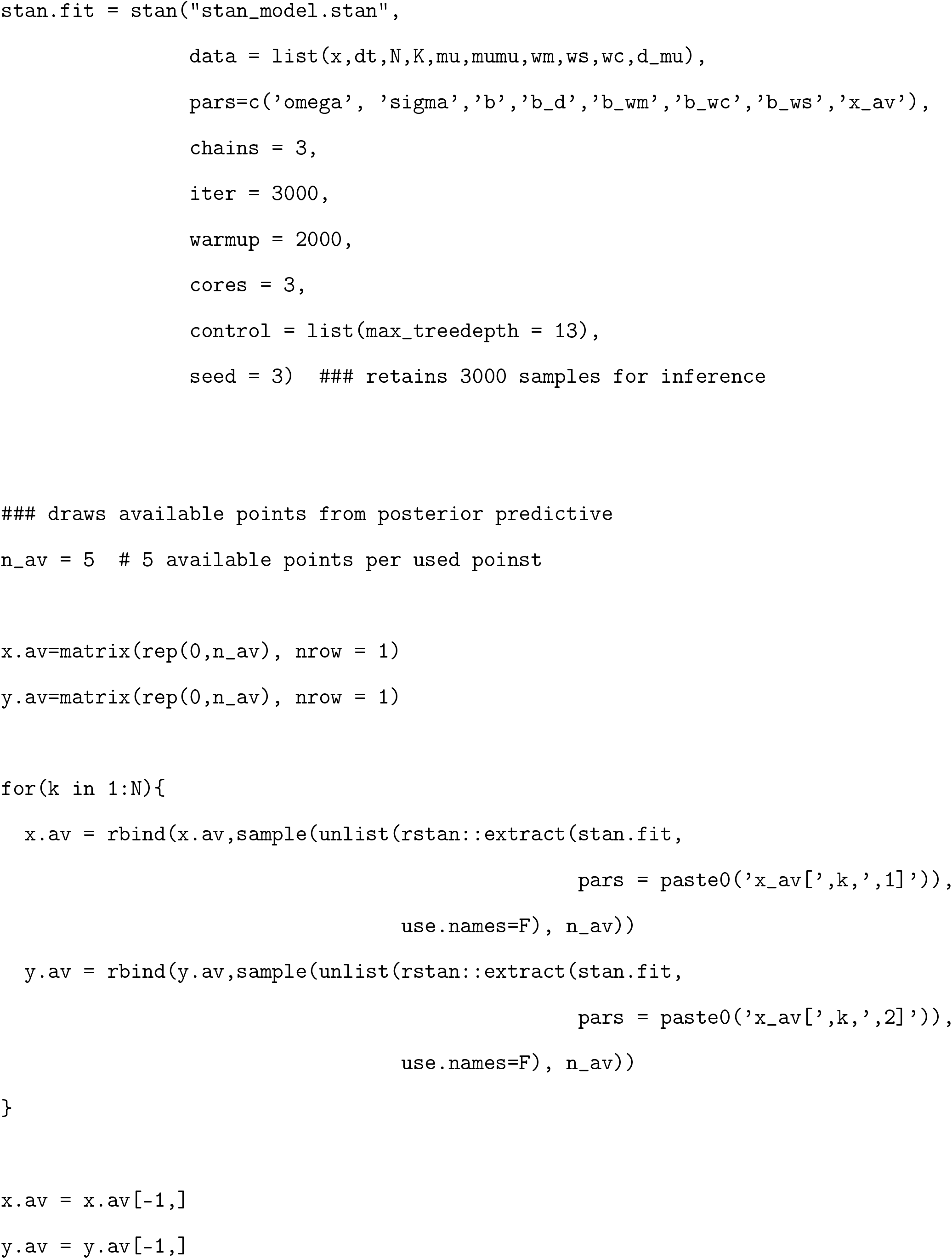
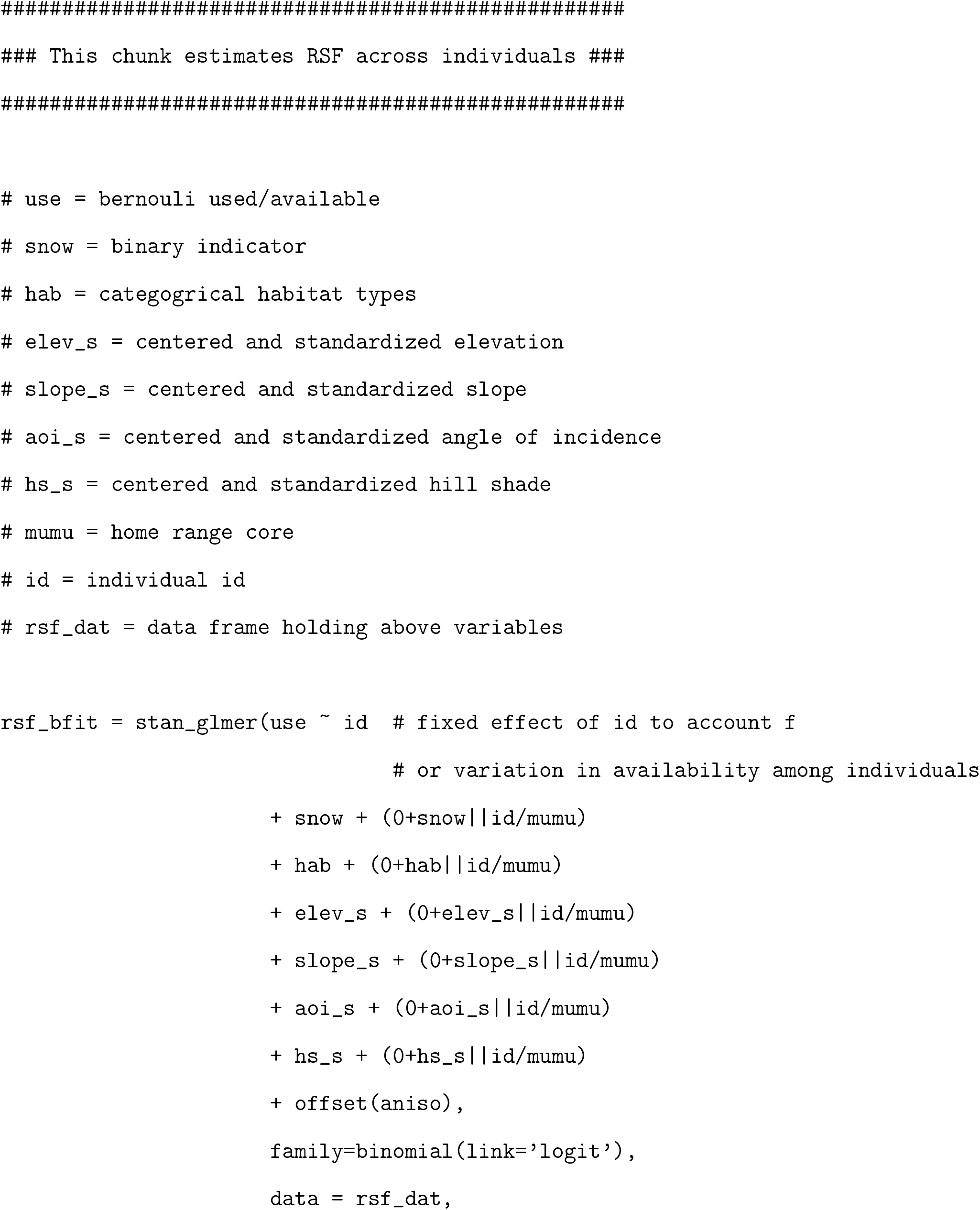
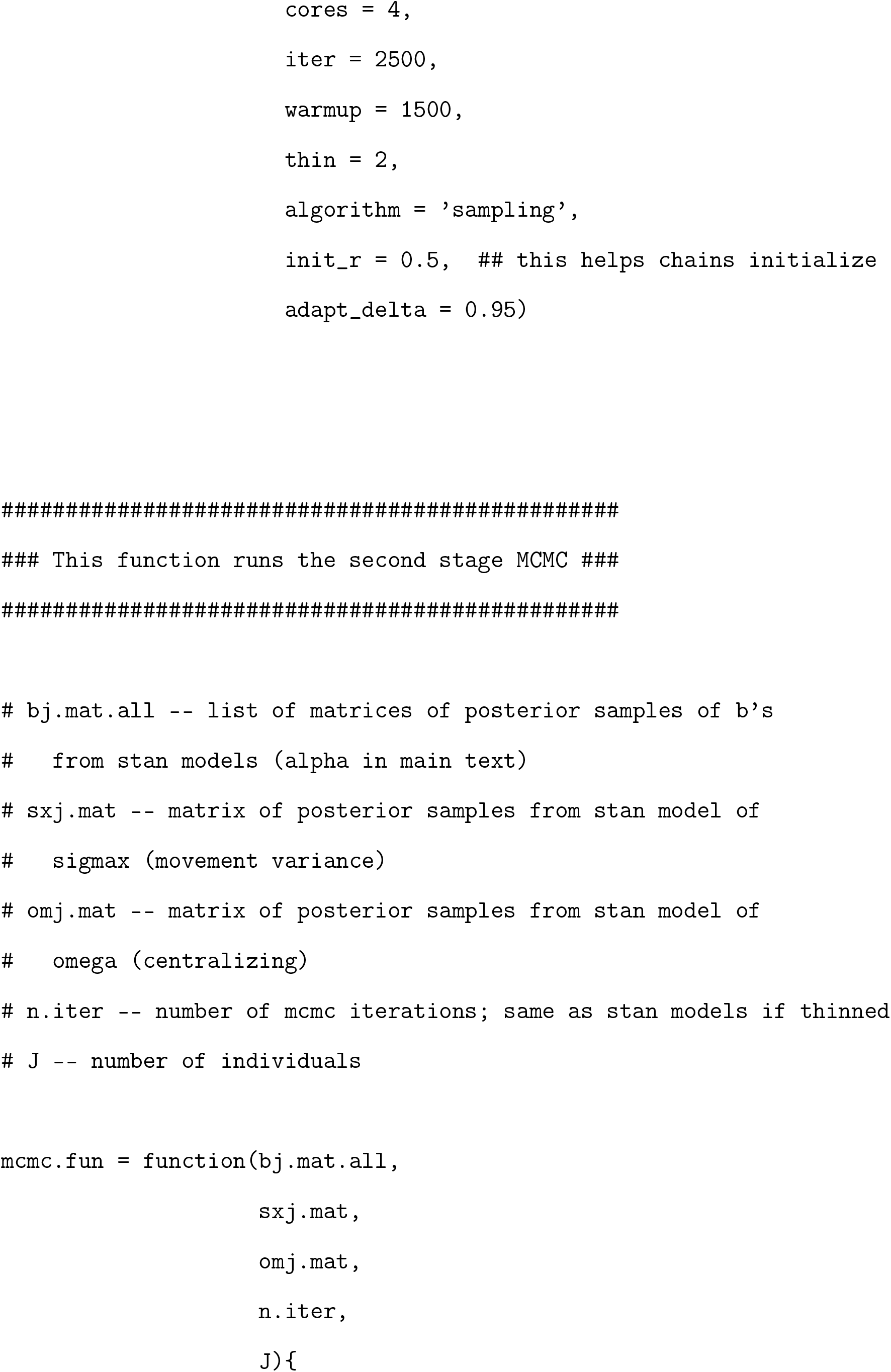
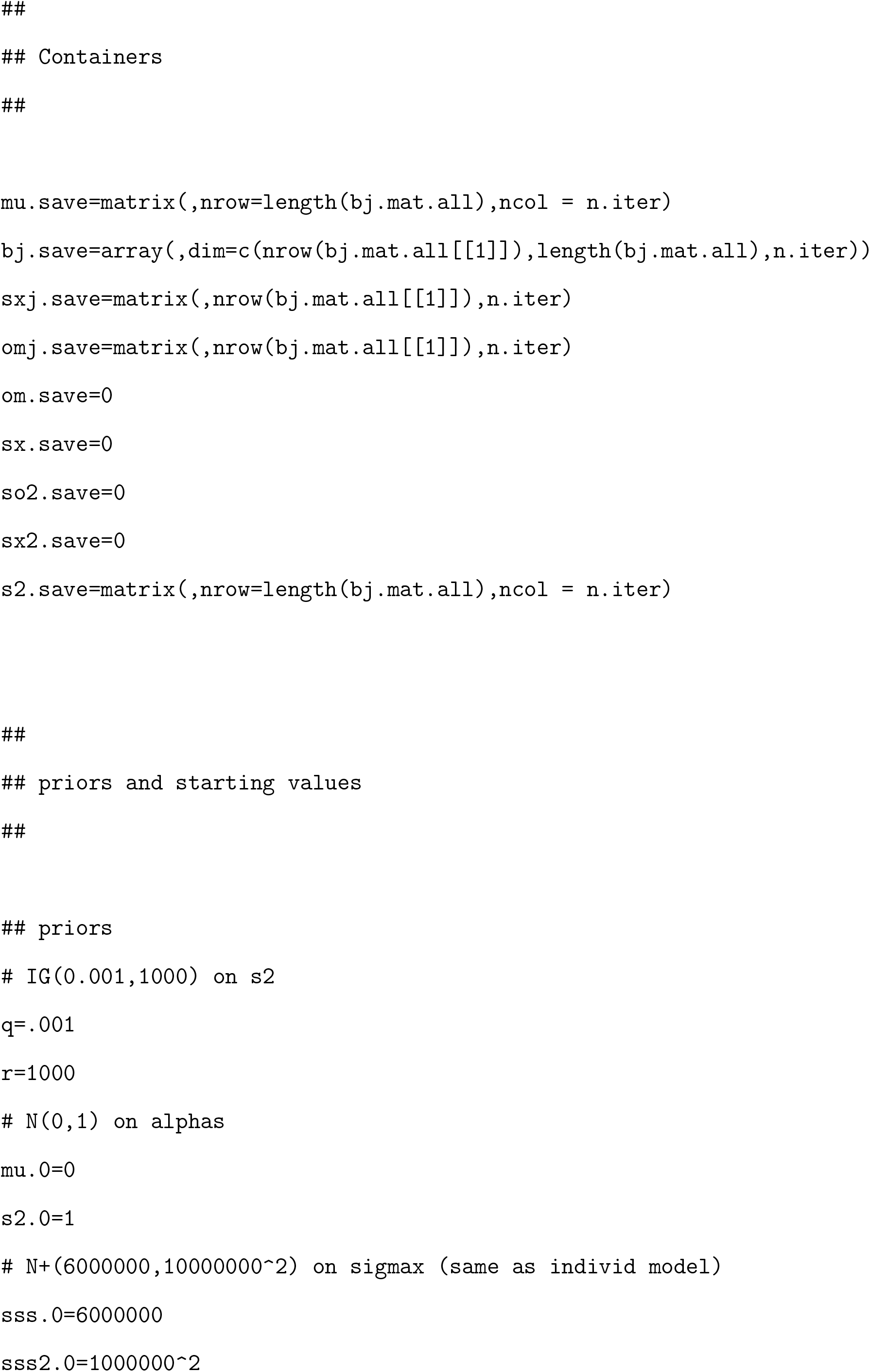
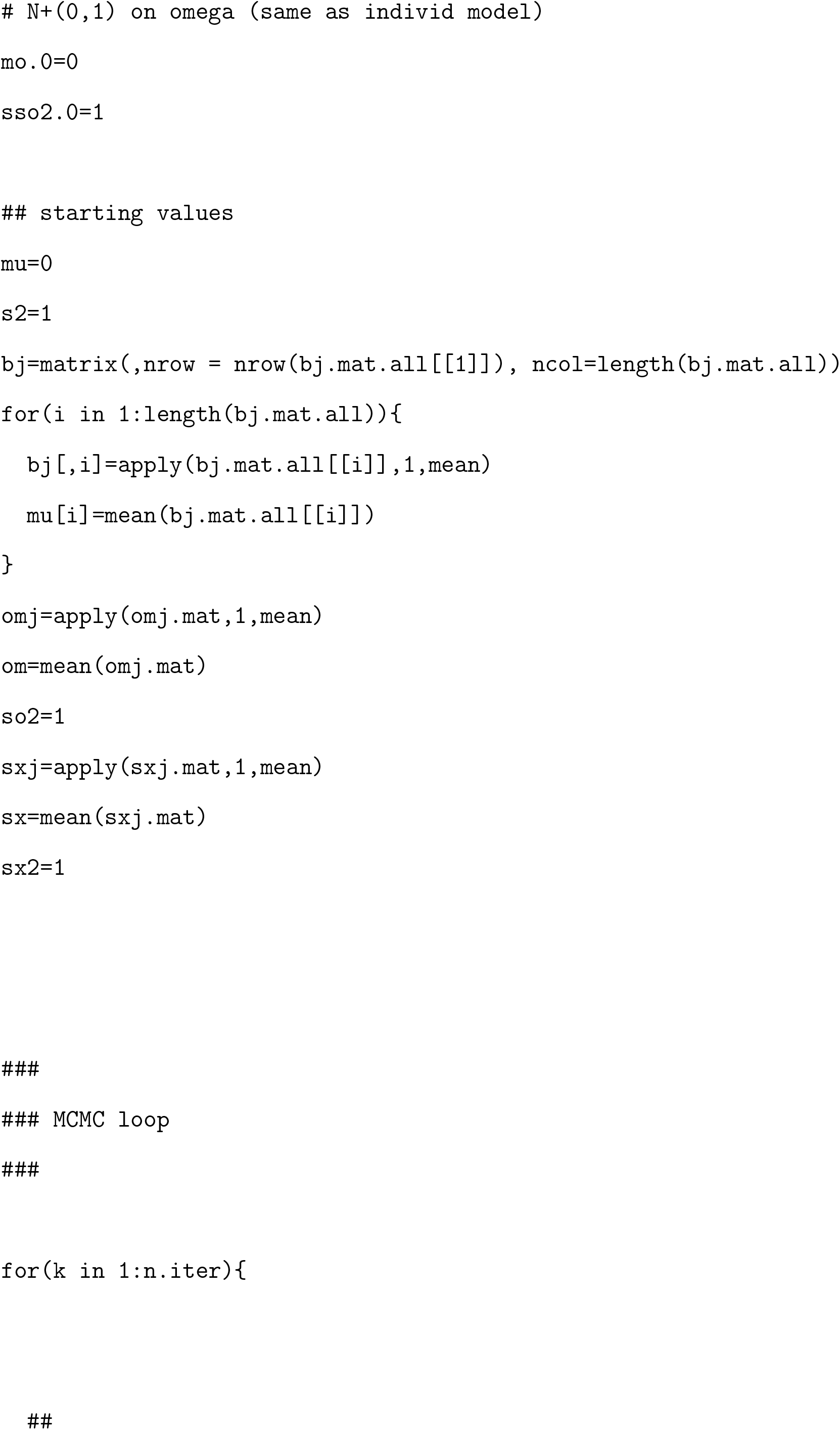
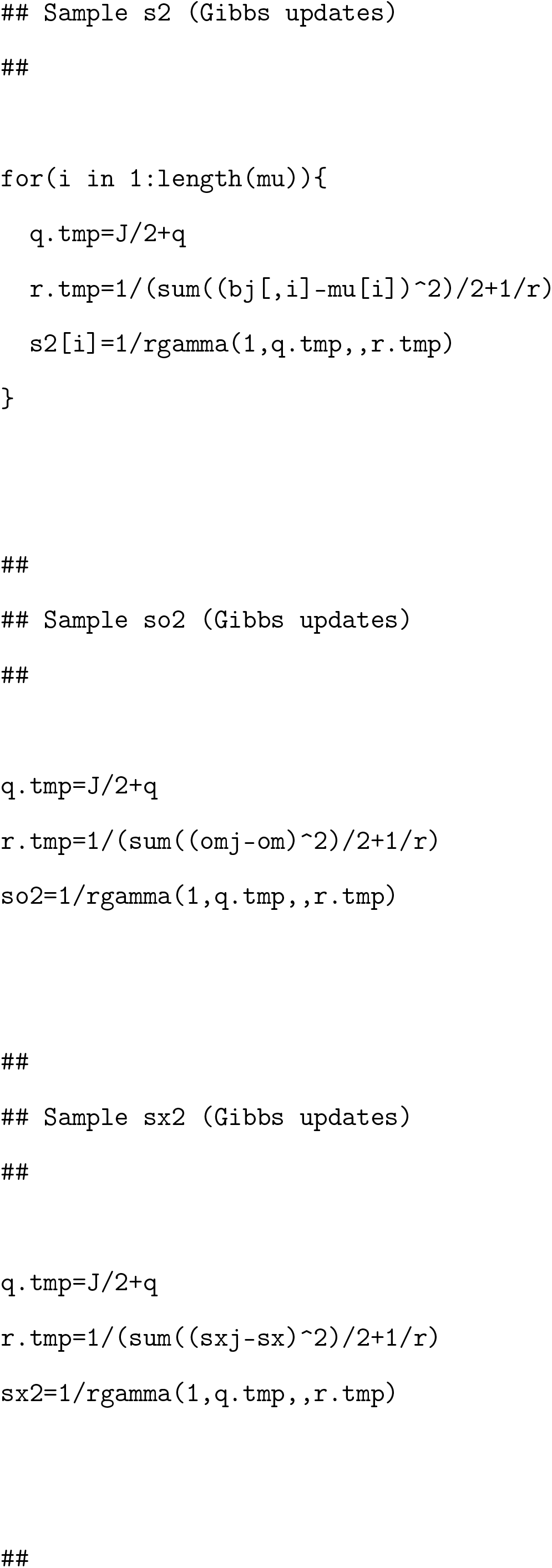
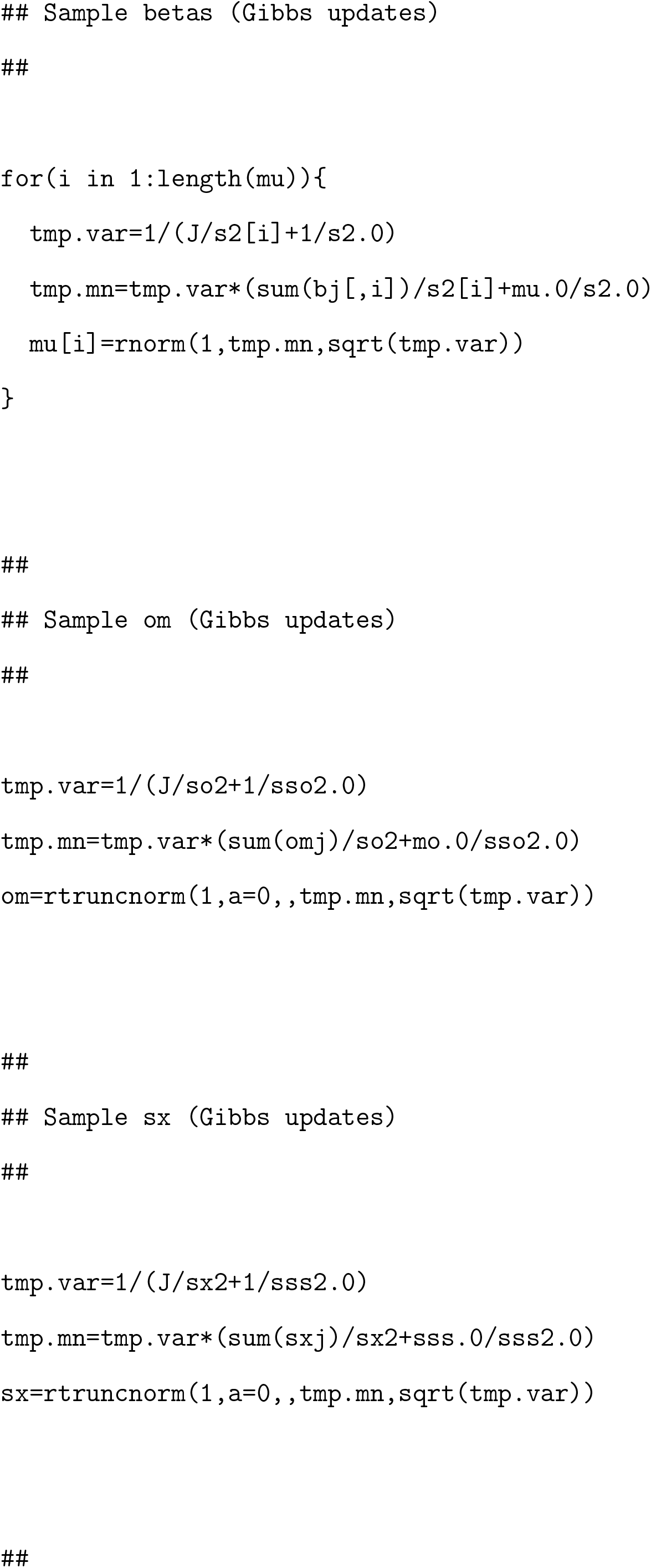
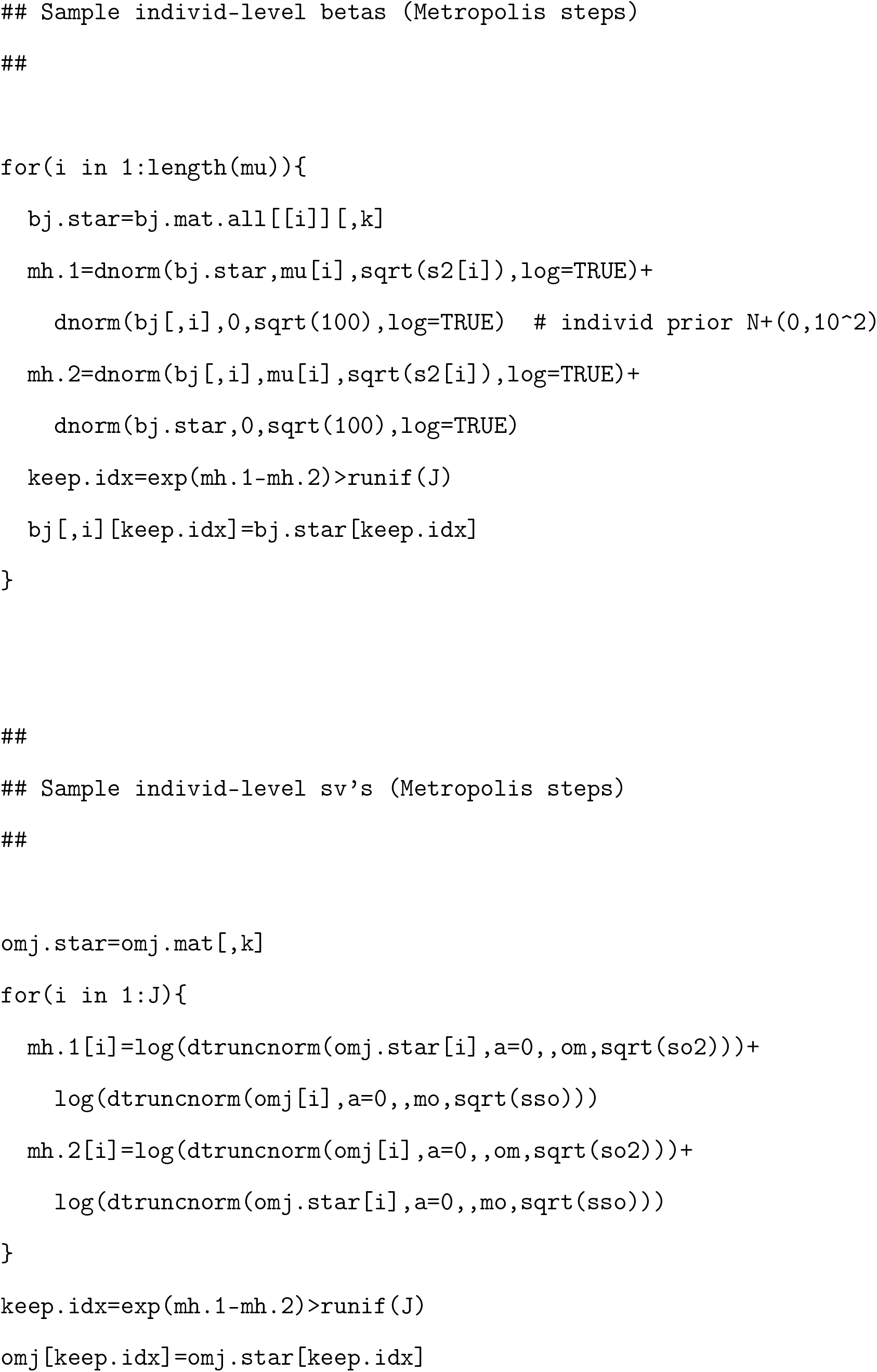
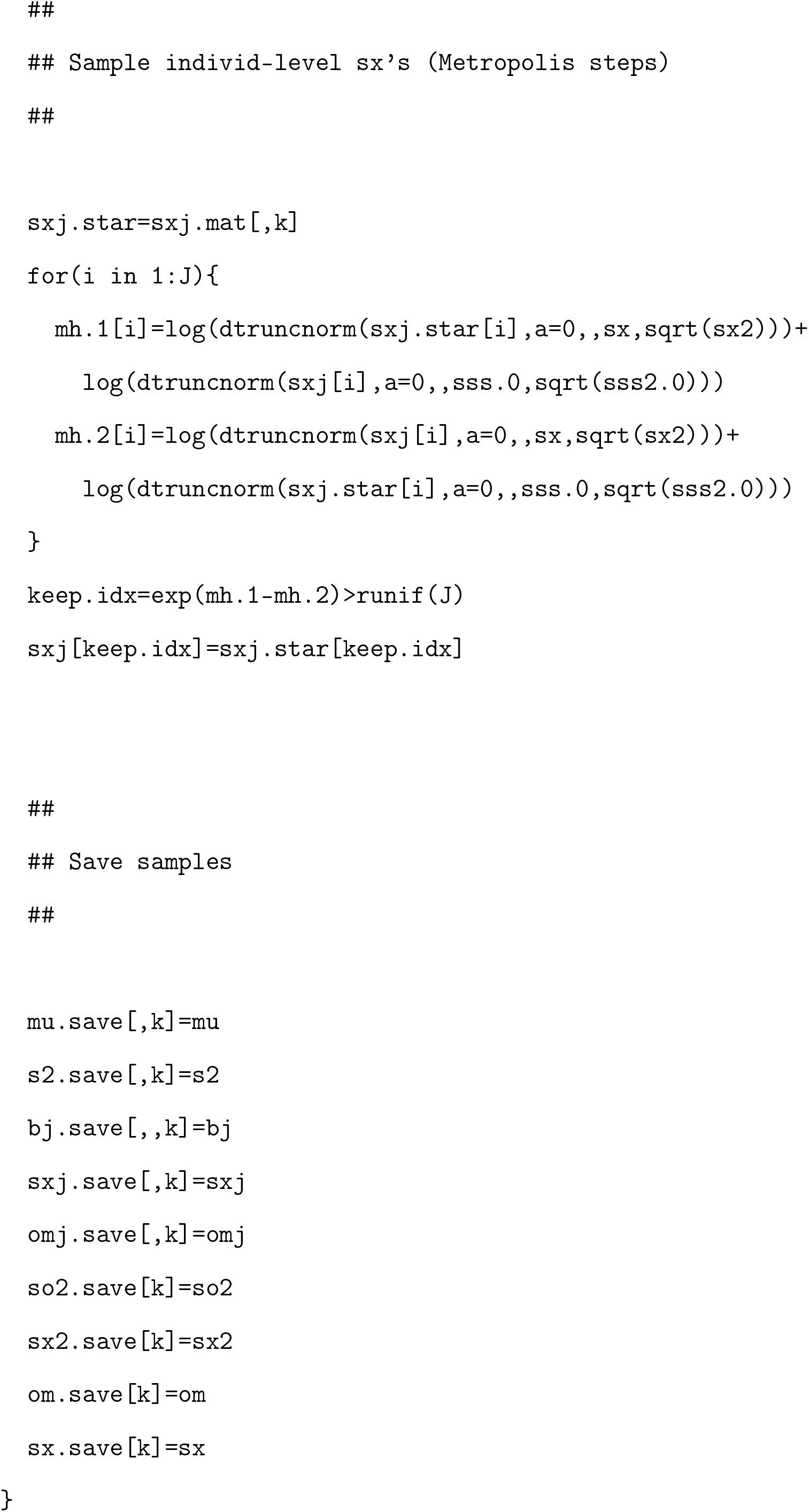
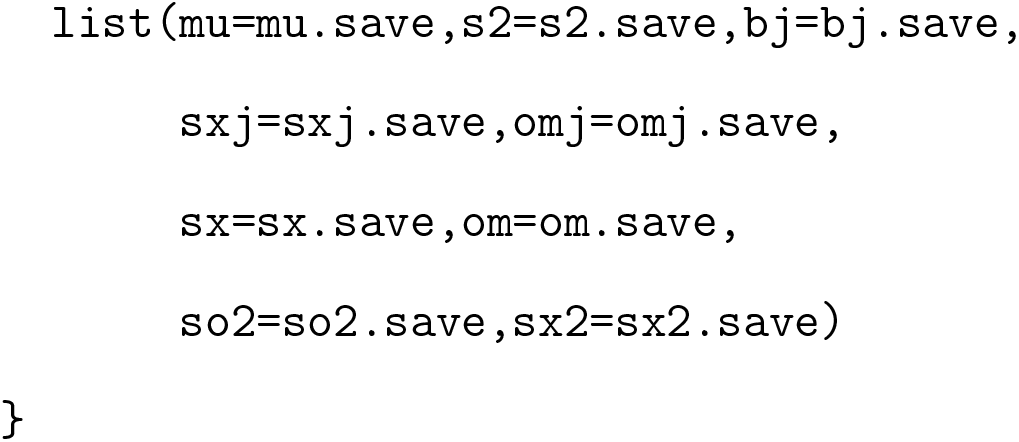

